# Meta-analysis and Open-source Database for In Vivo Brain Magnetic Resonance Spectroscopy in Health and Disease

**DOI:** 10.1101/2023.02.10.528046

**Authors:** Aaron T. Gudmundson, Annie Koo, Anna Virovka, Alyssa L. Amirault, Madelene Soo, Jocelyn H. Cho, Georg Oeltzschner, Richard A.E. Edden, Craig Stark

## Abstract

Proton (^1^H) Magnetic Resonance Spectroscopy (MRS) is a non-invasive tool capable of quantifying brain metabolite concentrations *in vivo*. Prioritization of standardization and accessibility in the field has led to the development of universal pulse sequences, methodological consensus recommendations, and the development of open-source analysis software packages. One on-going challenge is methodological validation with ground-truth data. As ground-truths are rarely available for *in vivo* measurements, data simulations have become an important tool. The diverse literature of metabolite measurements has made it challenging to define ranges to be used within simulations. Especially for the development of deep learning and machine learning algorithms, simulations must be able to produce accurate spectra capturing all the nuances of *in vivo* data. Therefore, we sought to determine the physiological ranges and relaxation rates of brain metabolites which can be used both in data simulations and as reference estimates. Using the Preferred Reporting Items for Systematic reviews and Meta-Analyses (PRISMA) guidelines, we’ve identified relevant MRS research articles and created an open-source database containing methods, results, and other article information as a resource. Using this database, expectation values and ranges for metabolite concentrations and T_2_ relaxation times are established based upon a meta-analyses of healthy and diseased brains.

## 1. Introduction

*In vivo* MRS can measure levels of metabolites in the brain non-invasively, allowing the abnormal biochemical and cellular processes of disease to be interrogated. The most prominent signals in the ^1^H spectrum are the methyl singlets associated with N-acetylaspartate/N-acetylaspartylglutamate (tNAA), creatine-containing compounds (tCr), and choline-containing compounds (tCho). Substantial multiplet contributions to the spectrum are also seen from myo-inositol (mI), glutamate (Glu), glutamine (Gln), gamma-aminobutyric acid (GABA), glutathione (GSH), and lactate (Lac). A handful of other metabolites can be quantified, including but not limited to: aspartate (Asp); ascorbate (Asc); scyllo-inositol (sI); serine (Ser); glycine (Gly); and taurine (Tau) [1–3]. For each of these metabolites, there exists a diffuse literature of measurements made using different methodologies in healthy controls and various populations of neurologic, psychiatric, and neurodevelopmental disease. Consensus on the physiological ranges for metabolite concentrations and relaxation values has yet to be determined.

Quantification of metabolite levels by MRS is challenging and a variety of methods are used to convert detected signal voltages into concentration-like measurements. These are all relative – that is, they rely upon the collection of a reference signal. Phantom-replacement [4] and synthetic referencing [5] are cumbersome and not widely used, so internal signal referencing predominates [6,7]. Among the potential reference signals, there is no clear and unambiguous ‘best’ option, each having advantages and disadvantages. Metabolite-metabolite referencing (most commonly to creatine) has the advantage of being simultaneously acquired and relatively unaffected by changing amounts of cerebrospinal fluid (CSF) within the measurement volume [8]. However, metabolite-water referencing is now the consensus-recommended approach, based upon the high SNR of the water signal and its role as the solvent [7,9,10]. Concentrations can be inferred from signal ratios and an assumption of the MR-visible water concentration, and can be expressed in molal (mol/kg solvent), molar (mol/dm^3^) or institutional units (i.u.) [7,9–11]. Correction for the varying water signal relaxation rates and visibilities in gray matter (GM), white matter (WM) and CSF is usually also performed on the basis of segmented structural images [12]. The relaxation of metabolite signals is usually corrected on the basis of literature reference values [12,13].

Generating realistic synthetic *in vivo* spectra is desirable for the development and validation of MRS quantification methods. Simulations that produce spectra that are fully representative of *in vivo* data, in terms of metabolite concentrations, macromolecular background, spectral baseline, artifacts and other nuances of MRS, will improve validation of classical methods and permit the development of deep learning techniques. Density matrix simulations based upon prior knowledge of metabolite chemical shifts and coupling constants [14–19] can generate metabolite basis spectra. However, deriving the metabolite component of a synthetic spectrum from simulated basis sets additionally requires specifying appropriate metabolite concentrations and lineshapes (combining relaxation behavior and field inhomogeneity). The International Society for Magnetic Resonance in Medicine (ISMRM) ‘Fitting Challenge’ was one of the first efforts to create realistic synthetic spectra to test the performance of different modeling software packages [20], specifying a single metabolite T_2_ value of 160 ms and, ‘normal ranges,’ for metabolite concentrations. While there have been a few disease-specific meta-analyses of MRS literature [21–25], there has not been a meta-analysis of the healthy and ‘control’ literature nor a cross-diagnosis comparison of the MRS literature. Therefore, in this manuscript we describe an open-source database which can be used to identify trends among the MRS literature and provide a meta-analysis to better inform future efforts to generate synthetic data that represent brain MRS in health and disease.

## 2. Methods

In the current study, we have developed a comprehensive open-source database that includes metabolite relaxation and concentration values. This collates the results of nearly 500 MRS papers, tabulating metabolite concentrations and relaxation rates for the healthy brain and a wide range of pathologies. Each entry also includes the publication information, experimental parameters, and data acquisition methods. To demonstrate the utility of this database, we performed three separate analyses: 1) an investigation into healthy brain metabolite concentrations; 2) a model of how these concentrations change in 25 clinical populations; and 3) a model to predict and account for variable metabolite T_2_ results.

### 2.1 Search Methods

In building the database, publications were identified to determine eligibility for inclusion according to Preferred Reporting Items for Systematic reviews and Meta-Analyses (PRISMA) guidelines [26,27]. Searches were conducted on PubMed, Web of Science, and Scopus databases. Separate searches (each search phrase is included in Supplementary Table 1) were carried out to specifically identify publications that either quantified metabolite concentrations or T_2_ relaxation times, herein referred to as the concentration study and relaxation studies, respectively. The original search for both was conducted in August of 2021. In order to include literature published throughout 2021, an additional follow-up search was conducted in March 2022. No limitation for publication date was specified for searches; only articles available in English were included. A PRISMA flowchart that reflects the process of building concentration and relaxation databases is shown in Figure 1.

**Figure 1:**
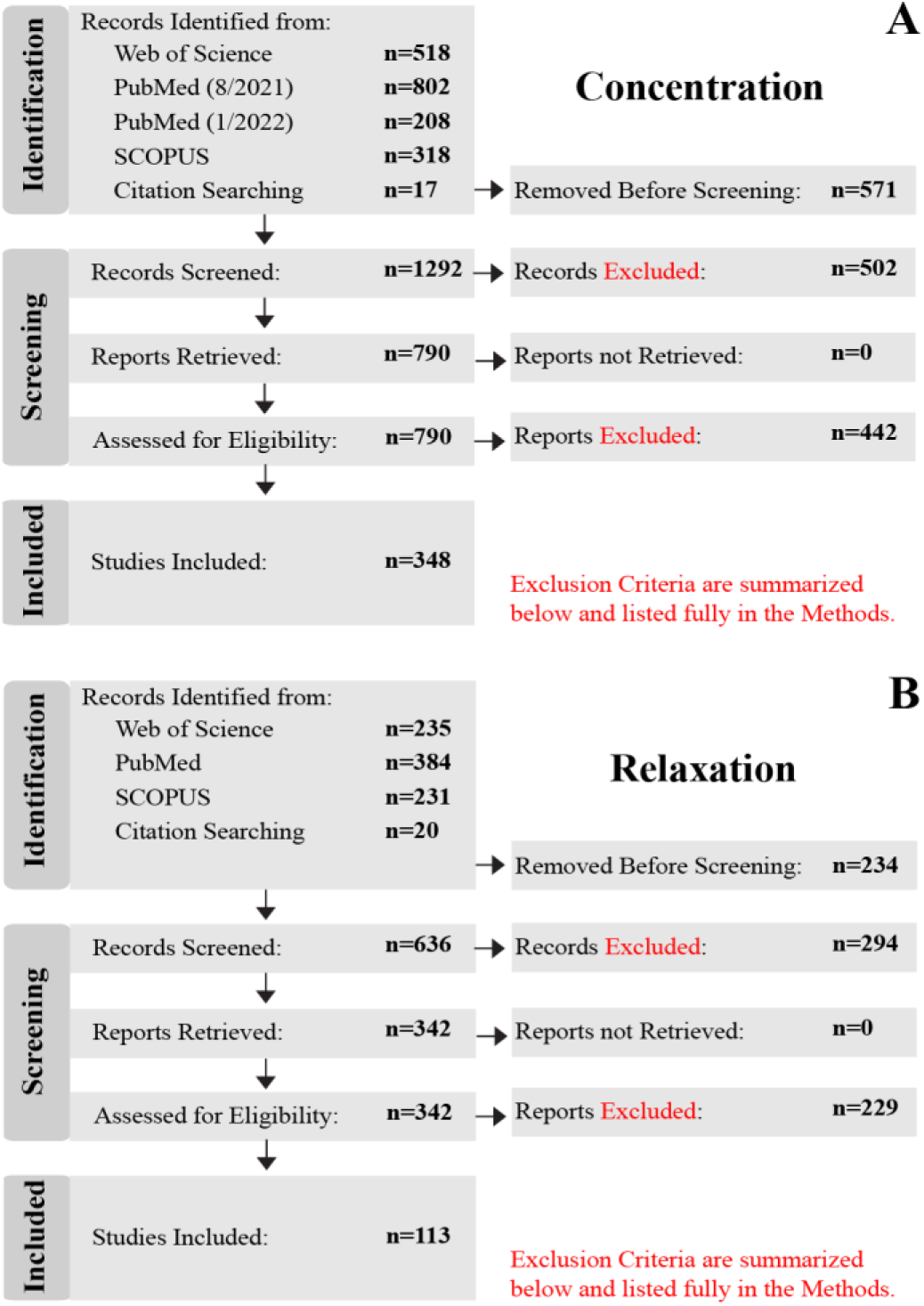
PRISMA flow charts that show the database selection and inclusion process of the **(A)** concentration and **(B)** T_2_ relaxation publications. Records “Removed Before Screening” were duplicates (identified from more than one database), reviews, meta-analyses, textbooks, or re-analyses. Conference abstracts were generally excluded, with exceptions made when information for a given metabolite/disease was scarce. “Records Excluded” were those identified as the wrong field of study, non-^1^H MRS, non-spectroscopy MR methods, non-brain regions, or animal studies (for Concentration study only). “Reports Excluded” during the “Assessed for Eligibility” did not include metabolite concentrations nor relaxation values.

For both the concentration and relaxation studies, only *in vivo* brain ^1^H-MRS data from primary sources were considered. Duplicate records (i.e., abstracts/titles) identified from more than one database, reviews, meta-analyses, re-analyses, and book chapters were excluded during the “Removed Before Screening” step of the “Identification” stage. Conference posters were generally also excluded at the “Removed Before Screening” step since they are not peer-reviewed (with exceptions made, where information was otherwise scarce). During the “Screening” stage, “Records Excluded” were those identified as the wrong field of study (e.g., NMR Spectroscopy for food science), X-nuclei (e.g., ^13^C, ^31^P, ^17^O, ^15^N, ^23^Na, etc.), non-spectroscopy MR methods (e.g., anatomical, functional, diffusion, etc.), or that did not study the brain. For the Concentration study, “Records Excluded” also included animal studies. All reports (i.e., research articles) were able to be retrieved for the remaining screened records. Finally, during the “Assessed for Eligibility” step, the “Reports Excluded” step reflects articles that did not include the mean and standard deviation for at least one metabolite concentration quantified in molar (moles/liter), molal (moles/g), or institutional units (i.u.), or referenced to total creatine (1/tCr), nor transverse relaxation times T_2_ or rates R_2_. Mean and standard deviation were calculated for reports listing median and quartile results, using the methods outlined in [28,29] to handle normal and skewed distributions, respectively. Distributions were classified as normal or skewed by comparing the upper and lower quartile-to-median ranges; if the range between the median and the lower quartile was similar to the range between and the median and the upper quartile (<50% difference), then the distribution was classified as normal, otherwise it was classified as skewed. Articles that presented values in the form of bar or scatter plots were included by manually determining mean and standard deviations with the assistance of an in-house Python software package that maps pixel values to figure axes. Authors were contacted by email ’ if they collected relevant data, but did not list their results; these included authors that only provided statistical results (e.g., t-statistic, p-value, etc.), non-standard units (e.g., arbitrary units), or normalized measurements (e.g., relative to baseline, z-scored, etc.).

Due to the high volume of articles (10,506) returned for the concentration study, articles were initially limited to 2018-2021. Where necessary, articles were retrieved from earlier years to ensure that three or more studies were included for less commonly studied clinical populations or difficult-to-measure metabolites (e.g., ascorbate) – this provided an abbreviated subset of 1,863 articles in the “Identification” stage. Of the original 1,863 articles, 571 articles were “Removed Before Screening” leaving 1,292 articles. After screening, 790 records remained and the corresponding report was retrieved. A total of 348 articles were determined to be eligible for inclusion in the database and analysis.

While this work aims to determine MRS features in the human brain, the relaxation study included all species as a handful of metabolites have not yet been well studied outside of animal models. A total of 870 articles were returned by the database searches during the “Identification” stage. Of the original 870 articles, 234 were “Removed Before Screening.” The remaining 636 records were “Screened” and 294 were removed in the “Records Excluded” step. 342 reports were then retrieved and assessed for eligibility. Finally, 113 articles remained and were included in the database and analyses.

Data were analyzed using in-house Python scripts that utilized *NumPy*, *Pandas*, *Scipy*, *Statsmodels*, *Matplotlib*, and *Scikit*-learn [30–35]. The weighted mean and 95% confidence intervals calculated within the healthy and clinical metabolite concentration meta-analyses used a combined effects model. Specifically, combined effects were determined using a Random Effects model [36] which can be advantageous for biological studies where a true value does not exist across studies (e.g., metabolite concentration varies from person to person). If a Random Effects model was not defined or there was not enough data (<8 studies), a Fixed Effect model was used [36] which can similarly identify common effects with less flexibility by assuming a singular true value. Weighting across studies, both for combined effects and meta-regression, used the inverse variance weighting scheme [37] to penalize high-variance studies. While all data are present in the database, meta-analyses were only carried out when 3 or more studies were available for a particular metabolite, group, or field strength.

### 2.2. Metabolite Concentrations in Healthy Populations

Studies that investigated healthy individuals or had healthy control groups were used to determine metabolite concentration ranges in healthy populations. Of the 350 studies included, 259 studies investigated a healthy population or included a healthy control group (26% of studies included no healthy subjects). Subjects were classified into early life (<2 years of age), adolescent (5-14 years of age), young adult (18-45 years of age) and aged adult (>50 years of age). These age ranges allowed for the greatest number of studies to be included in each of the categories while leaving a gap (e.g., 46-49 years of age) to set groups apart. There were 8 [38–44], 19 [45–63], 199 [54,64–258], and 45 [81,97,142,152,156,159,194,196,225,244,259–288] studies within the four age categories (early life, adolescent, young adult, aged), respectively. To determine the concentration ranges, values were separated by metabolite and units (i.u./mM and 1/tCr) reported. Finally, a combined effects model [36] was used to compute the mean and 95% confidence interval (as seen in Figure 2.

**Figure 2:**
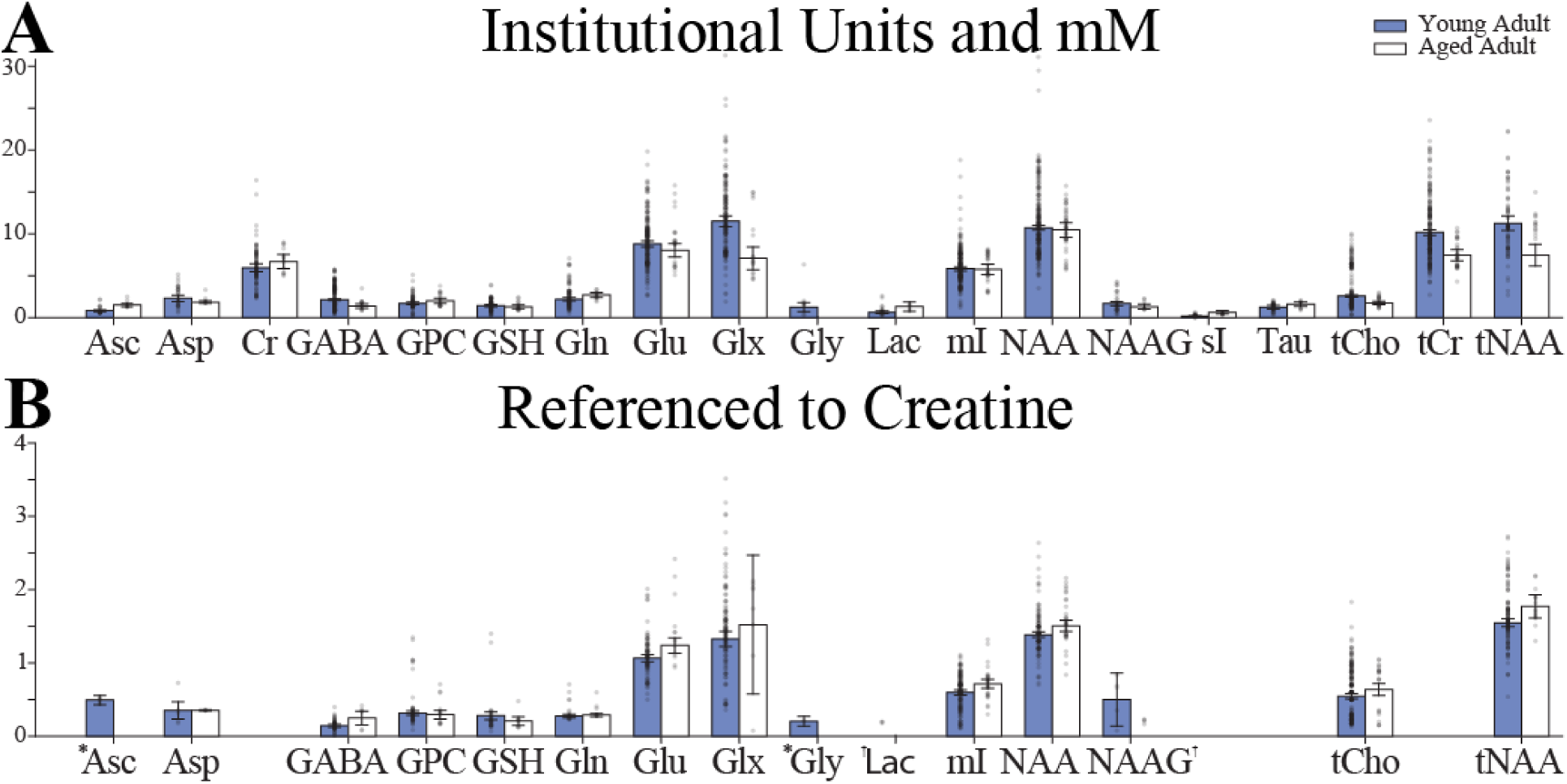
Brain metabolite concentrations in younger (18-45 years, in blue) and older (>50 years, in white) healthy adults from studies that reported results as: **(A)** Molar, molal, and Institutional Units; **(B)** Creatine-referenced. An * indicates the use of a Fixed Effects Model rather than a Random Effects Model. A † indicates a combined effects model was not defined.

### 2.3. Metabolite Concentrations in Clinical Populations

Studies that investigated clinical groups and included a healthy control group were included in the clinical population analysis. There were 180 publications [38–43,49–52,54–57,59,62–64,68,69,71,73–75,77,78,80,82,84,87,91,92,94–98,100,101,103,104,106,107,110,111,113,116–118,120,124,128,129,136–142,144,146,148–151,153,155,160–165,170,173,174,176,178,179,181–183,185–189,191,192,196–200,202–205,207,209,210,213,218,220–222,224,227,229,230,232,235,237,240,242,243,246–248,250–255,257–261,263,264,267–273,276,277,279–313] consisting of 25 unique clinical groups. To determine the concentration ranges, values were separated by metabolite and units reported. Each clinical population was then modeled as a linear change relative to their respective control group by using the ‘ratio of means’ method [314,315]. A value of 1.0 would indicate no difference between the clinical and control groups. Finally, a combined effects model [36] was used to compute the mean and 95% confidence interval (as seen in Figure 3).

**Figure 3:**
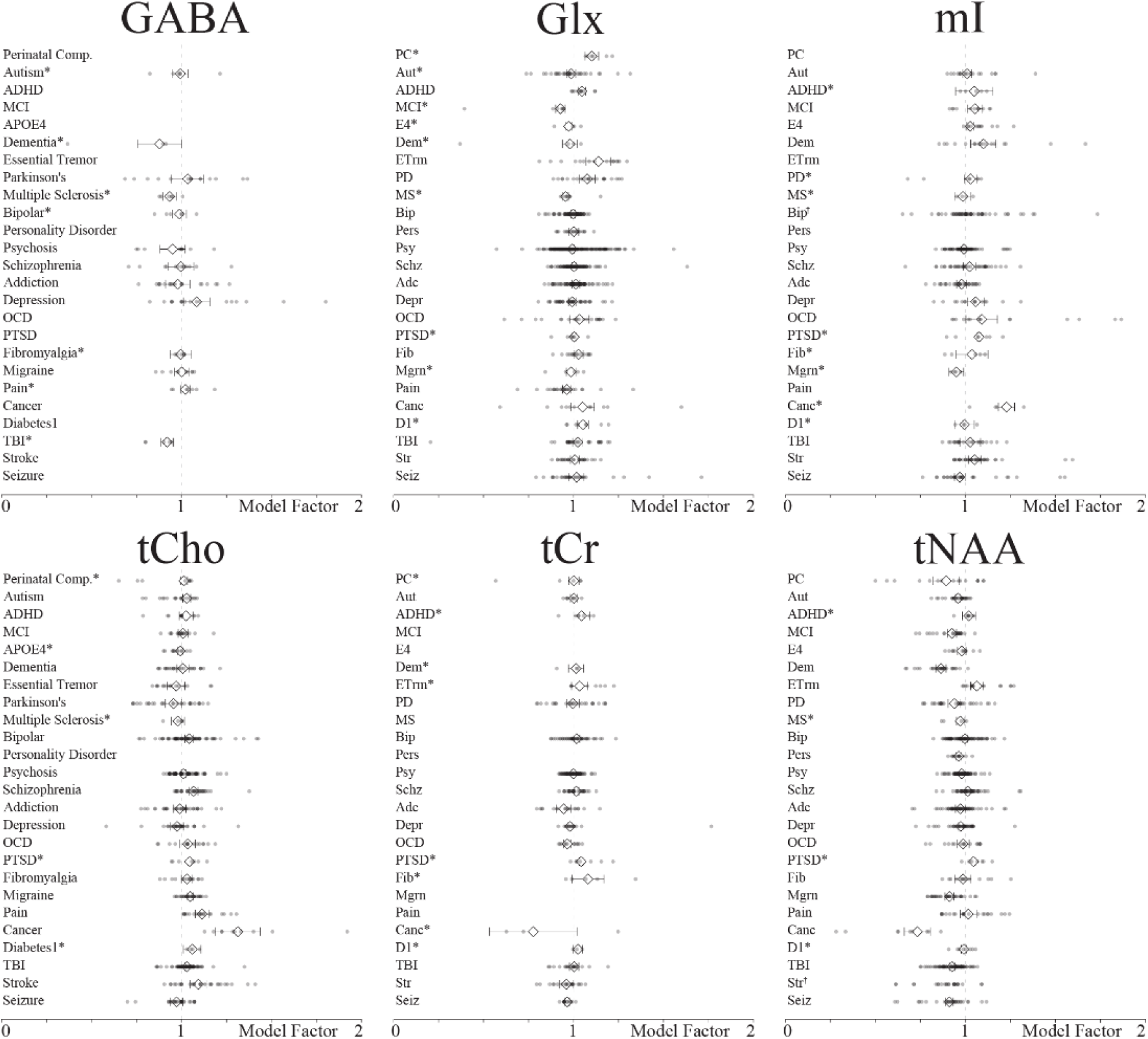
The six most commonly investigated metabolite concentrations modeled in diseased Populations. Data from metabolite and metabolite complexes are combined (e.g., Cr and tCr, Glu and Glx). An * by the group classification indicates the use of a Fixed Effects Model rather than a Random Effects Model. A † indicates a combined effects model was not defined. PC = perinatal complications; Aut = autism; ADHD = attention-deficit/hyper activity; MCI = mild cognitive impairment; E4 apolipoprotein 4 carriers; Dem = dementia; Etrm = essential tremor; PD = Parkinson’s disease; MS = multiple sclerosis; Bip = bipolar; Pers = personality disorder; Psy = psychosis; Schz = schizophrenia; Adc = addiction; Depr = depression; OCD = obsessive compulsive disorder; PTSD = post-traumatic stress disorder; Fib = fibromyalgia; Mgrn = migraine; Pain = chronic pain; Canc = cancer; D1 = type 1 diabetes; TBI = traumatic brain injury; Str = stroke; Seiz = seizure disorder.

### 2.4. T_2_ Meta-regression Model

Studies that investigated healthy subjects or included a healthy control group were included in the T_2_ relaxation analysis. Of the 113 included studies, 76 studies [3,13,316–389] were included in the analysis. All the studies’ results were separated by metabolite for the analysis to produce 629 values. Next, a multiple meta-regression was employed with 6 input variables: 1) metabolite; 2) field strength; 3) localization pulse sequence; 4) T_2_ filter, 5) tissue type; and 6) subject species. Metabolite was a categorical variable that included 14 metabolites, with some of them further differentiated by moiety (Asp, tCr CH_2_, Cr CH_3_, GABA, Gln, Glu, Gly, tCho, GSH, Lac, mI, NAA CH_3_, NAAG, Tau). Field strength was a continuous variable from 1.5 T through 14.1 T. Localization pulse sequence was a categorical variable that included Point Resolved Spectroscopy (PRESS), Stimulated Echo Acquisition Mode (STEAM), or either Localization by Adiabatic Selective Refocusing (LASER) or semi-LASER (sLASER). ‘T_2_ filter’ was a categorical variable indicating whether the data were collected with a Carr-Purcell Meiboom-Gill (CPMG) multi-echo sequence or not. Tissue type was a categorical variable which was characterized as GM (voxel composition >80% GM), WM (voxel composition >80% WM), or mixed (all other cases). Subject species was a categorical variable that specified human or not human. The output was a continuous T_2_ value in milliseconds. Continuous variables were scaled between 0 and 1. Categorical variables were dummy coded creating for use within the regression model. The model was iteratively re-run leaving one datapoint out each time for prediction (i.e., 629 individual leave-one-out regression models were run).

## 3. Results

### 3.1. Database

The database currently contains 461 publications with each entry containing the publication information, experiment details, parameters of the data acquisition, and the mean and standard deviation of the results. A complete list of the information available from each entry in the database is given in Table 1. We used the PRISMA guidelines to ensure an unbiased and wide-reaching approach was taken to identify and screen publications. The database is open-source and available online at https://github.com/agudmundson/mrs-database.

**Table 1.** Information available for entries in the database.

|  |  |
| --- | --- |
| **Citation:** | **Voxel:** |
| Name in Database | Dimensions (x, y, z) |
| Publication Year | Volume |
| Author(s) | Anatomical Region |
| Journal Volume | Hemisphere |
| Title | Tissue Fractions (Mean/Standard Deviation) |
| Digital Object Identifier |  |
|  | **Acquisition:** |
| **Study Populations:** | Localization Sequence |
| Study Index | Water Suppression |
| Population | Acquisition Bandwidth |
| Control Group | Number of Datapoints |
| Treatment or Conditions | Number of Transients |
| Visit or Session Number | Repetition Time (TR) |
| Total Number of Subjects | Echo Time (TE) |
| Number of Subjects Analyzed | Inversion Time (TI) |
| Number of Male Subjects | T<sub>2</sub> Filter |
| Number of Female Subjects |  |
| Age (Mean/Standard Deviation) | **Analysis:** |
|  | Preprocessing Software |
| **Hardware:** | Fitting/Quantification Software |
| Scanner Manufacturer | Segmentation Software |
| Scanner Model | Partial Volume Correction |
| Magnetic Field Strength | Relaxation Correction |

### 3.2. Healthy Metabolite Concentrations

The physiological ranges of brain metabolites were determined within the each of the four age categories for both i.u./mM and 1/tCr. The resulting weighted mean and 95% confidence intervals for young and aged adult concentrations, for both i.u./mM and 1/tCr, are shown in Figure 2. The weighted mean, 95% confidence intervals, and other summary statistics for healthy infant, adolescent, young adult, and aged populations are available at https://github.com/agudmundson/mrs-database.

### 3.3. Clinical Metabolite Concentrations in pathological conditions

While clinical studies that did not include a control group were included in the database, the main focus was on studies that had direct comparisons, to minimize confounds involving technical variations among studies. Rather than computing effect sizes, linear changes were used to be directly interpretable to generate concentrations for future simulations. Figure 3 depicts levels of commonly investigated metabolites measured in diseased populations. The mean linear change, 95% confidence intervals, and other summary statistics for each metabolite in the 25 clinical populations is available at https://github.com/agudmundson/mrs-database.

### 3.4. T_2_ relaxation

The iterative leave-one-out models achieved a median adjusted R^2^ of 0.782 (Q1 = .7817; Q3 = 0.7819). Predictions for these models yielded a median error of 26.61 ms (Q1 = 12. 06 ms; Q3 = 54.66 ms) with 16.23% error (Q1 = 7.51%; Q3 = 27.29%). Figure 4 shows the actual value plotted with the marker size representing the weight within the model and the meta-regression model for 3 of the most common metabolites, NAA, Cho, Cr. The full model is available at https://github.com/agudmundson/mrs-database.

**Figure 4:**
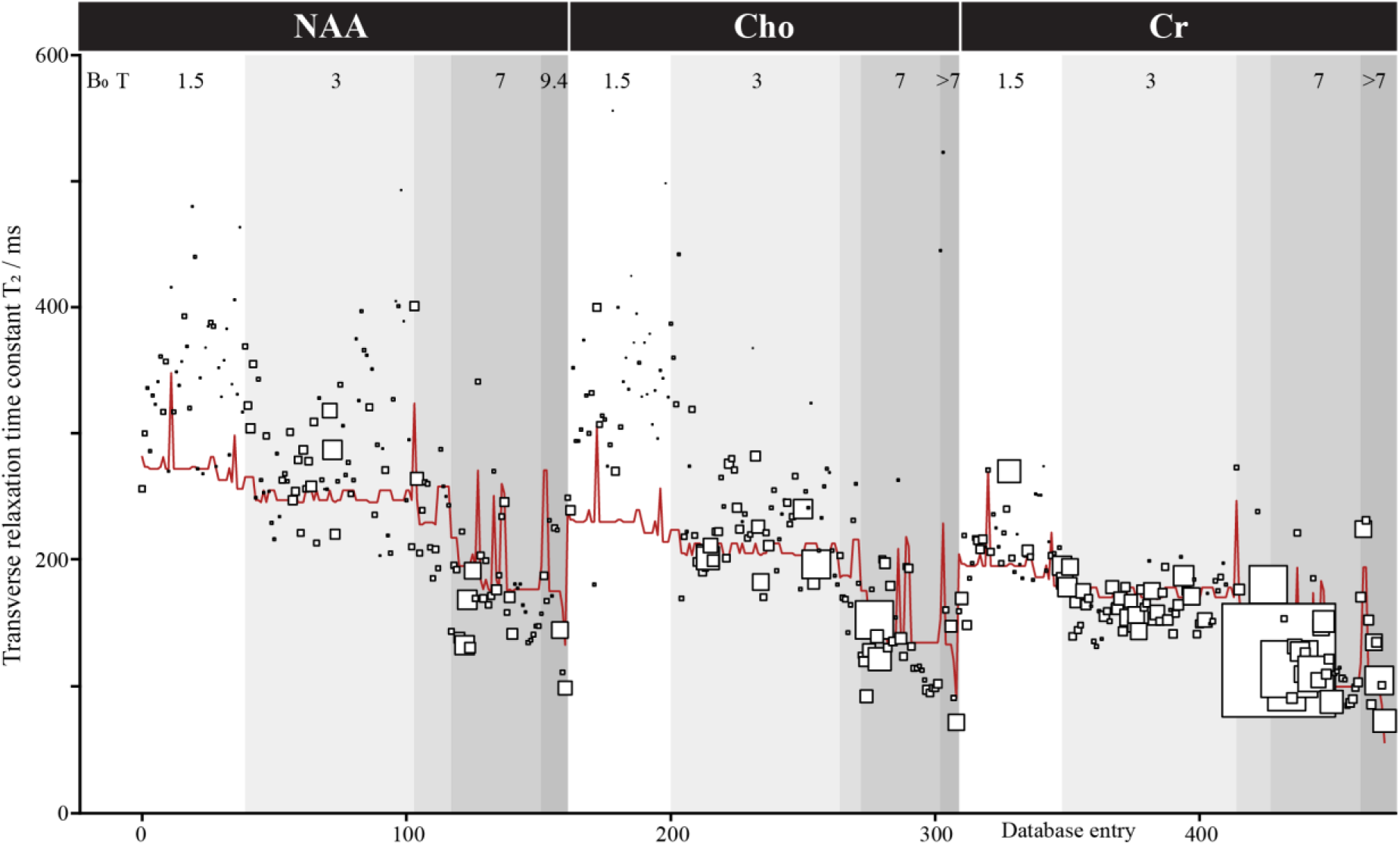
Transverse relaxation time meta-analysis. Only results for NAA, Cho, and Cr are shown for ease of visualization, but a total of 629 values for 14 metabolites were included in the database and modeled. Metabolite, field strength, localization, T_2_ filter, species, and tissue type were included as factors in the model. Database entries are sorted here by these factors in that order. Each study is represented by a square of size reflecting the modeling weight (based on the inverse of variance). The red line shows the model.

## 4. Discussion

### 4.1 Open-source Database

Using a systematic approach, we provide the first database for MRS results and corresponding methods. As this database is freely available through the cloud-based website GitHub, new entries can be continually added and existing entries can be updated with more information through collaborative efforts. This database is valuable for quickly identifying trends as results across multiple studies can be interrogated. As with the meta-analyses performed here, future analyses may interrogate brain region, software, or other methodological decisions.

### 4.2 Physiological Ranges of Brain Metabolites in the Healthy Adults

The primary goal of this meta-analysis was to summarize levels of MRS-accessible metabolites with a large data mining and unification approach. This was not the first effort to provide typical concentration values or ranges – physiological ranges of metabolites have been proposed previously for the healthy brain using data from multiple species [390,391]. Here, a comprehensive approach was taken to unify measures across hundreds of human studies and appropriately weight them to establish the physiological ranges of 19 brain metabolites and metabolite-complexes. The focus here on recent publications (<5 years old) biased the analysis toward data quantified using more current and advanced methodologies. Reassuringly, many values here reflect similar ranges to those previously proposed [20,390,391].

Methodological heterogeneity within the MRS literature certainly limits any quantitative meta-analysis. The extent to which such effects negatively impact this analysis vary. For example, the clinical effects are all quantified within-study (i.e., each study is characterized in terms of a fractional group difference), so first-order effects associated with different tissue corrections shifting the mean concentration values are not a concern; however, second-order effects (where less valid corrections might lead to a group-bias) are still a concern. The main concentration analysis does conflate data quantified with a variety of methods. The variance observed in the meta-analysis results thus includes measurement variance as well as methodological variance within the literature. Where the combined/random effects models ‘compare’ categories with biased sampling among the methods represented, this approach may be misled by methodological biases.

The metabolic profile provided here represents progress towards effective and accurate simulation of realistic synthetic data. The development of data analysis methodologies is limited by a lack of ground truths – methodological performance is usually assessed in terms of modeling uncertainty (CRLB) or within- or between-subject variance (standard deviation). Notably, these metrics do not reflect a true measurement error, tending to ignore measurement bias and conflate sources of variance. Ultimately, synthetic data that accurately represent all features of *in vivo* data allow comprehensive evaluation of sources of variance and bias in MRS methods. Beyond validation of traditional analysis methods, such synthetic data are integral to developing deep learning and machine learning algorithms for MRS data analysis and quantification.

### 4.3. Physiological Ranges of Brain Metabolites in Clinical Populations

Here, a linear model demonstrating the relationship between healthy and clinical populations was presented. Results for the six most frequently quantified metabolites can be seen in Figure 3. As far as we know, this is the first study to provide a basis to determine physiological and pathological ranges of brain metabolites in such a wide array of clinical populations. Many of the cohort effects summarized, and those highlighted in Figure 3, agree with previous systematic reviews and disease-specific meta-analyses [21–25,392]. For example, our analysis reproduced the widely recognized elevated choline in tumors [22] and elevated mI and decreased NAA in aging and Alzheimer’s Disease [21,392–394]. Neurometabolic changes in tNAA and tCho also appear to have some value in discriminating between clinical syndromes with similar symptomology, such as Parkinson’s Disease and Essential Tremor [395–397]. It is notable that, although tCr is often used as an internal reference, it is not markedly stable across the disease populations, and can show changes across aging [398,399]. By synthesizing meta-analytic information across a range of disorders, this resource may allow the development of future tools to discriminate between clinical conditions.

### 4.4. Multiple Meta-Regression to Explain Heterogeneity of Metabolite T_2_ Relaxation Results

T_2_ relaxation is an important aspect of *in vivo* MRS data and should be carefully considered when simulating data. Unfortunately, apart from the 3 most common methyl singlets (i.e., tNAA, tCr, tCho), T_2_ ranges have not been well established. This can be seen as most relaxation-corrected absolute quantification methods rely on a small handful of references and must make approximations for tissue differences, pulse sequence effects, or even for metabolites that have not been studied for the given acquisition protocol. The goal of this analysis was to produce a model that could provide metabolite T_2_ ranges for simulation. To do this, we leveraged data from multiple metabolites across different species that were measured using a variety of acquisition schemes. While results between studies can be seen to have a high degree of variability, the multiple meta-regression model was able to account for a large degree of the variance. The model included 6 variables: 1) metabolite, 2) field strength, 3) localization pulse sequence, 4) T_2_ filter, 5) tissue type, and 6) subject species. Following a leave-one-out validation approach, nearly 80% of the variance could be attributed to the 6 factors. The major factors that explain variance in T_2_ are field strength, with shorter T_2_ at higher field; metabolite, with Cr having shorter T_2_ than Cho and especially NAA; species, with longer T_2_ in rodents; and T_2_-filter (although CPMG filters are only used in a minority of studies). The error in prediction was low, with approximately 25% of the prediction errors less than 10 ms, 50% of prediction errors less than 25 ms, and nearly 75% of prediction errors under 50 ms. High prediction errors came primarily from a small subset of papers that appear to represent outliers in the dataset suggesting predictions may provide reliable estimates when simulating understudied metabolites. We did not attempt to quantify ‘study quality’ as a potential weighting factor, other than through cohort size. The main factor that is not included in the model (although addressed to some degree by the ‘tissue factor’) is brain region of measurements, where iron-rich regions are known to show shorter T_2_s [400–402]. It will also be important to measure T_2_ data in clinical populations and across the lifespan to further solidify the existing body of literature.

The context of generating the metabolite components of synthetic spectra, which this meta-analysis builds toward, requires metabolite basis functions, metabolite amplitudes, as well as metabolite linewidths. By surveying the metabolite T_2_ literature, we sought to better understand the ‘pure-T_2_’ of metabolite linewidths in the absence of local field inhomogeneity. One model of *in vivo* lineshapes is to assume T_2_ as the Lorentzian linewidth component and to assign inhomogeneity broadening to a Gaussian linewidth component – the data compiled here can inform such a model. It is also appropriate to consider transverse relaxation weighting of metabolite amplitudes when assembling synthetic spectra “acquired” at typical TEs.

## 5. Conclusion

Here, we provide a new database containing brain metabolite results from nearly 500 MRS publications. This database is freely available online where users can view and contribute their own data. Moving forward, this database can function as a community resource allowing deeper interrogation and understanding of how acquisition protocol, software analysis tools, brain region, population, etc. impact and/or bias results.

Using the database, we have determined physiological ranges of 19 brain metabolites and metabolite-complexes across the lifespan of healthy individuals. We further modeled disease effects relative to healthy controls to allow for determining concentration ranges for 25 psychiatric and neurologic diseases. Finally, we have performed a meta-regression to determine appropriate ranges for T_2_ in MRS simulations. The determined ranges will be invaluable for informing the generation of synthetic data for evaluating analysis tools and deep learning datasets. Additionally, these ranges may serve as a reference to clinical researchers that are unaware of the expected values for a given metabolite or may be considering how MRS can fit within their study design.

## Abbreviations

^1^H: proton
2-HG: 2-hydroxyglutarate
Adc: addiction
ADHD: attention-deficit/hyper activity
Asc: ascorbate
Asp: aspartate
Aut: autism
Bip: bipolar
Canc: cancer
Cho: choline-containing compounds
CPMG: Carr-Purcell Meiboom-Gill
Cr: creatine
CRLB: Cramer-Rao lower bounds
CSF: cerebrospinal fluid
D1: type 1 diabetes
Dem: dementia
Dep: depression
E4: apolipoprotein 4 carriers
Etrm: Essential Tremor
Fib: fibromyalgia
GABA: gamma-aminobutyric acid
Gln: glutamine
Glu: glutamate
Glx: sum of glutamate and glutamine
Gly: glycine
GM: gray matter
GPC: glycerophosphocholine
ISMRM: international society for magnetic resonance in medicine
Lac: lactate
LASER: localization by adiabatic selective refocusing
MCI: mild cognitive impairment
MEGA: Mescher-Garwood
mI: myo-inositol
Mig: migraine
MRS: magnetic resonance spectroscopy
MS: multiple sclerosis
NAA: N-acetylaspartate
NAAG: N**-**acetyl-aspartyl-glutamate
OCD: obsessive compulsive disorder
Pain: chronic pain
PC: perinatal Complications
PCho: phosphocholine
PCr: phosphocreatine
PD: Parkinson’s disease
PE: phosphoethanolamine
Pers: personality disorder
PRISMA: preferred reporting Items for systematic reviews and meta-analyses
PRESS: point resolved spectroscopy
Psy: psychosis
PTSD: post-traumatic stress disorder
Schz: schizophrenia
Seiz: seizure disorder
Ser: serine
sI: scyllo-inositol
sLASER: semi-adiabatic localization by adiabatic selective refocusing
STEAM: stimulated echo acquisition mode
SNR: signal-to-noise ratio
Str: stroke
T_2_: spin-spin relaxation time
Tau: taurine
TBI: traumatic brain injury
tCho: sum of choline-containing metabolites
tCr: sum of creatine and phosphocreatine
tNAA: sum of N-acetyl-aspartate and N-acetyl-aspartyl-glutamate
TE: echo-time
TI: inversion time
TM: mixing time
TR: repetition time
WM: white matter

## References

[1] A. Henning, Proton and multinuclear magnetic resonance spectroscopy in the human brain at ultra-high field strength: A review, Neuroimage. 168 (2018) 181–198. https://doi.org/10.1016/j.neuroimage.2017.07.017.

[2] D.K. Deelchand, I. Iltis, P.-G. Henry, Improved quantification precision of human brain short echo-time 1 H magnetic resonance spectroscopy at high magnetic field: A simulation study, Magn. Reson. Med. 72 (2014) 20–25. https://doi.org/10.1002/mrm.24892.

[3] J. Pfeuffer, I. Tkáč, S.W. Provencher, R. Gruetter, Toward an in Vivo Neurochemical Profile: Quantification of 18 Metabolites in Short-Echo-Time 1H NMR Spectra of the Rat Brain, J. Magn. Reson. 141 (1999) 104–120. https://doi.org/10.1006/jmre.1999.1895.

[4] B.J. Soher, P.C.M. vanZijl, J.H. Duyn, P.B. Barker, Quantitative proton MR spectroscopic imaging of the human brain, Magn. Reson. Med. 35 (1996) 356–363. https://doi.org/10.1002/mrm.1910350313.

[5] L. Barantin, A. Le Pape, S. Akoka, A new method for absolute quantitation MRS metabolites, Magn. Reson. Med. 38 (1997) 179–182. https://doi.org/10.1002/mrm.1910380203.

[6] G. Öz, D.K. Deelchand, J.P. Wijnen, V. Mlynárik, L. Xin, R. Mekle, R. Noeske, T.W.J. Scheenen, I. Tkáč, O. Andronesi, P.B. Barker, R. Bartha, A. Berrington, V. Boer, C. Cudalbu, U.E. Emir, T. Ernst, A. Fillmer, A. Heerschap, P.G. Henry, R.E. Hurd, J.M. Joers, C. Juchem, H.E. Kan, D.W.J. Klomp, R. Kreis, K. Landheer, S. Mangia, M. Marjańska, J. Near, E.M. Ratai, I. Ronen, J. Slotboom, B.J. Soher, M. Terpstra, J. Valette, M. Van der Graaf, M. Wilson, Advanced single voxel 1H magnetic resonance spectroscopy techniques in humans: Experts’ consensus recommendations, NMR Biomed. 34 (2021) 1–18. https://doi.org/10.1002/nbm.4236.

[7] J. Near, A.D. Harris, C. Juchem, R. Kreis, M. Marjańska, G. Öz, J. Slotboom, M. Wilson, C. Gasparovic, Preprocessing, analysis and quantification in single-voxel magnetic resonance spectroscopy: experts’ consensus recommendations, NMR Biomed. 34 (2021) 1–23. https://doi.org/10.1002/nbm.4257.

[8] B.S.Y. Li, H. Wang, O. Gonen, Metabolite ratios to assumed stable creatine level may confound the quantification of proton brain MR spectroscopy, Magn. Reson. Imaging. 21 (2003) 923–928. https://doi.org/10.1016/S0730-725X(03)00181-4.

[9] P.B. Barker, B.J. Soher, S.J. Blackband, J.C. Chatham, V.P. Mathews, R.N. Bryan, Quantitation of proton NMR spectra of the human brain using tissue water as an internal concentration reference, NMR Biomed. 6 (1993) 89–94. https://doi.org/10.1002/nbm.1940060114.

[10] R. Kreis, T. Ernst, B.D. Ross, Absolute Quantitation of Water and Metabolites in the Human Brain. II. Metabolite Concentrations, J. Magn. Reson. Ser. B. 102 (1993) 9–19. https://doi.org/10.1006/jmrb.1993.1056.

[11] C. Gasparovic, T. Song, D. Devier, H.J. Bockholt, A. Caprihan, P.G. Mullins, S. Posse, R.E. Jung, L.A. Morrison, Use of tissue water as a concentration reference for proton spectroscopic imaging, Magn. Reson. Med. 55 (2006) 1219–1226. https://doi.org/10.1002/mrm.20901.

[12] C. Gasparovic, H. Chen, P.G. Mullins, Errors in 1H-MRS estimates of brain metabolite concentrations caused by failing to take into account tissue-specific signal relaxation, NMR Biomed. 31 (2018) 1–9. https://doi.org/10.1002/nbm.3914.

[13] V. Mlynárik, S. Gruber, E. Moser, Proton T 1 and T 2 relaxation times of human brain metabolites at 3 Tesla, NMR Biomed. 14 (2001) 325–331. https://doi.org/10.1002/nbm.713.

[14] R. Simpson, G.A. Devenyi, P. Jezzard, T.J. Hennessy, J. Near, Advanced processing and simulation of MRS data using the FID appliance FID-A —An open source, MATLAB-based toolkit, Magn. Reson. Med. 77 (2017) 23–33. https://doi.org/10.1002/mrm.26091.

[15] S.A. Smith, T.O. Levante, B.H. Meier, R.R. Ernst, Computer Simulations in Magnetic Resonance. An Object-Oriented Programming Approach, J. Magn. Reson. Ser. A. 106 (1994) 75–105. https://doi.org/10.1006/jmra.1994.1008.

[16] Z. Starčuk, J. Starčuková, Quantum-mechanical simulations for in vivo MR spectroscopy: Principles and possibilities demonstrated with the program NMRScopeB, Anal. Biochem. 529 (2017) 79–97. https://doi.org/10.1016/j.ab.2016.10.007.

[17] H.J. Hogben, M. Krzystyniak, G.T.P. Charnock, P.J. Hore, I. Kuprov, Spinach - A software library for simulation of spin dynamics in large spin systems, J. Magn. Reson. 208 (2011) 179–194. https://doi.org/10.1016/j.jmr.2010.11.008.

[18] S.C.N. Hui, M.G. Saleh, H.J. Zöllner, G. Oeltzschner, H. Fan, Y. Li, Y. Song, H. Jiang, J. Near, H. Lu, S. Mori, R.A.E. Edden, MRSCloud: A cloud-based MRS tool for basis set simulation, Magn. Reson. Med. 88 (2022) 1994–2004. https://doi.org/10.1002/mrm.29370.

[19] K. Landheer, K.M. Swanberg, C. Juchem, Magnetic resonance Spectrum simulator (MARSS), a novel software package for fast and computationally efficient basis set simulation, NMR Biomed. 34 (2021) 1–13. https://doi.org/10.1002/nbm.4129.

[20] M. Marjańska, D.K. Deelchand, R. Kreis, J.R. Alger, P.J. Bolan, T. Borbath, F. Boumezbeur, C.C. Fernandes, E. Coello, B.H. Nagraja, M. Považan, H. Ratiney, D. Sima, J. Starčuková, B.J. Soher, M. Wilson, J.J.A. van Asten, Results and interpretation of a fitting challenge for MR spectroscopy set up by the MRS study group of ISMRM, Magn. Reson. Med. 87 (2022) 11–32. https://doi.org/10.1002/mrm.28942.

[21] T. Song, X. Song, C. Zhu, R. Patrick, M. Skurla, I. Santangelo, M. Green, D. Harper, B. Ren, B.P. Forester, D. Öngür, F. Du, Mitochondrial dysfunction, oxidative stress, neuroinflammation, and metabolic alterations in the progression of Alzheimer’s disease: A meta-analysis of in vivo magnetic resonance spectroscopy studies, Ageing Res. Rev. 72 (2021) 101503. https://doi.org/10.1016/j.arr.2021.101503.

[22] Q. Wang, H. Zhang, J.S. Zhang, C. Wu, W.J. Zhu, F.Y. Li, X.L. Chen, B.N. Xu, The diagnostic performance of magnetic resonance spectroscopy in differentiating high-from low-grade gliomas: A systematic review and meta-analysis, Eur. Radiol. 26 (2016) 2670–2684. https://doi.org/10.1007/s00330-015-4046-z.

[23] A. Marsman, M.P. van den Heuvel, D.W.J. Klomp, R.S. Kahn, P.R. Luijten, H.E. Hulshoff Pol, Glutamate in Schizophrenia: A Focused Review and Meta-Analysis of 1H-MRS Studies, Schizophr. Bull. 39 (2013) 120–129. https://doi.org/10.1093/schbul/sbr069.

[24] K.M. Chitty, J. Lagopoulos, R.S.C. Lee, I.B. Hickie, D.F. Hermens, A systematic review and meta-analysis of proton magnetic resonance spectroscopy and mismatch negativity in bipolar disorder, Eur. Neuropsychopharmacol. 23 (2013) 1348–1363. https://doi.org/10.1016/j.euroneuro.2013.07.007.

[25] M.V. Vidor, A.C. Panzenhagen, A.R. Martins, R.B. Cupertino, C.E. Bandeira, F.A. Picon, B.S. da Silva, E.S. Vitola, L.A. Rohde, D.L. Rovaris, C.H.D. Bau, E.H. Grevet, Emerging findings of glutamate–glutamine imbalance in the medial prefrontal cortex in attention deficit/hyperactivity disorder: systematic review and meta-analysis of spectroscopy studies, Eur. Arch. Psychiatry Clin. Neurosci. 272 (2022) 1395–1411. https://doi.org/10.1007/s00406-022-01397-6.

[26] D. Moher, A. Liberati, J. Tetzlaff, D.G. Altman, Preferred reporting items for systematic reviews and meta-analyses: The PRISMA statement, BMJ. 339 (2009) 332–336. https://doi.org/10.1136/bmj.b2535.

[27] M.J. Page, D. Moher, P.M. Bossuyt, I. Boutron, T.C. Hoffmann, C.D. Mulrow, L. Shamseer, J.M. Tetzlaff, E.A. Akl, S.E. Brennan, R. Chou, J. Glanville, J.M. Grimshaw, A. Hróbjartsson, M.M. Lalu, T. Li, E.W. Loder, E. Mayo-Wilson, S. Mcdonald, L.A. Mcguinness, L.A. Stewart, J. Thomas, A.C. Tricco, V.A. Welch, P. Whiting, J.E. Mckenzie, PRISMA 2020 explanation and elaboration: Updated guidance and exemplars for reporting systematic reviews, BMJ. 372 (2021). https://doi.org/10.1136/bmj.n160.

[28] T. Greco, G. Biondi-Zoccai, M. Gemma, C. Guérin, A. Zangrillo, G. Landoni, How to impute study-specific standard deviations in meta-analyses of skewed continuous endpoints?, World J. Meta-Analysis. 3 (2015) 215. https://doi.org/10.13105/wjma.v3.i5.215.

[29] X. Wan, W. Wang, J. Liu, T. Tong, Estimating the sample mean and standard deviation from the sample size, median, range and/or interquartile range, BMC Med. Res. Methodol. 14 (2014) 1–13. https://doi.org/10.1186/1471-2288-14-135.

[30] C.R. Harris, K.J. Millman, S.J. van der Walt, R. Gommers, P. Virtanen, D. Cournapeau, E. Wieser, J. Taylor, S. Berg, N.J. Smith, R. Kern, M. Picus, S. Hoyer, M.H. van Kerkwijk, M. Brett, A. Haldane, J.F. del Río, M. Wiebe, P. Peterson, P. Gérard-Marchant, K. Sheppard, T. Reddy, W. Weckesser, H. Abbasi, C. Gohlke, T.E. Oliphant, Array programming with NumPy, Nature. 585 (2020) 357–362. https://doi.org/10.1038/s41586-020-2649-2.

[31] J.D. Hunter, Matplotlib: A 2D Graphics Environment, Comput. Sci. Eng. 9 (2007) 90–95. https://doi.org/10.1109/MCSE.2007.55.

[32] W. McKinney, Data Structures for Statistical Computing in Python, Proc. 9th Python Sci. Conf. 1 (2010) 56–61. https://doi.org/10.25080/majora-92bf1922-00a.

[33] F. Pedregosa, G. Varoquaux, A. Gramfort, V. Michel, B. Thirion, O. Grisel, M. Blondel, P. Prettenhofer, R. Weiss, V. Dubourg, J. Vanderplas, A. Passos, D. Cournapeau, M. Brucher, M. Perrot, E. Duchesnay, Scikit-learn: Machine Learning in {P}ython, J. Mach. Learn. Res. 12 (2011) 2825–2830.

[34] S. Seabold, J. Perktold, Statsmodels: Econometric and Statistical Modeling with Python, Proc. 9th Python Sci. Conf. (2010) 92–96. https://doi.org/10.25080/majora-92bf1922-011.

[35] P. Virtanen, R. Gommers, T.E. Oliphant, M. Haberland, T. Reddy, D. Cournapeau, E. Burovski, P. Peterson, W. Weckesser, J. Bright, S.J. van der Walt, M. Brett, J. Wilson, K.J. Millman, N. Mayorov, A.R.J. Nelson, E. Jones, R. Kern, E. Larson, C.J. Carey, İ. Polat, Y. Feng, E.W. Moore, J. VanderPlas, D. Laxalde, J. Perktold, R. Cimrman, I. Henriksen, E.A. Quintero, C.R. Harris, A.M. Archibald, A.H. Ribeiro, F. Pedregosa, P. van Mulbregt, A. Vijaykumar, A. Pietro Bardelli, A. Rothberg, A. Hilboll, A. Kloeckner, A. Scopatz, A. Lee, A. Rokem, C.N. Woods, C. Fulton, C. Masson, C. Häggström, C. Fitzgerald, D.A. Nicholson, D.R. Hagen, D. V. Pasechnik, E. Olivetti, E. Martin, E. Wieser, F. Silva, F. Lenders, F. Wilhelm, G. Young, G.A. Price, G.L. Ingold, G.E. Allen, G.R. Lee, H. Audren, I. Probst, J.P. Dietrich, J. Silterra, J.T. Webber, J. Slavič, J. Nothman, J. Buchner, J. Kulick, J.L. Schönberger, J.V. de Miranda Cardoso, J. Reimer, J. Harrington, J.L.C. Rodríguez, J. Nunez-Iglesias, J. Kuczynski, K. Tritz, M. Thoma, M. Newville, M. Kümmerer, M. Bolingbroke, M. Tartre, M. Pak, N.J. Smith, N. Nowaczyk, N. Shebanov, O. Pavlyk, P.A. Brodtkorb, P. Lee, R.T. McGibbon, R. Feldbauer, S. Lewis, S. Tygier, S. Sievert, S. Vigna, S. Peterson, S. More, T. Pudlik, T. Oshima, T.J. Pingel, T.P. Robitaille, T. Spura, T.R. Jones, T. Cera, T. Leslie, T. Zito, T. Krauss, U. Upadhyay, Y.O. Halchenko, Y. Vázquez-Baeza, SciPy 1.0: fundamental algorithms for scientific computing in Python, Nat. Methods. 17 (2020) 261–272. https://doi.org/10.1038/s41592-019-0686-2.

[36] M. Borenstein, L. V. Hedges, J.P.T. Higgins, H.R. Rothstein, Introduction to Meta-Analysis, John Wiley & Sons, Ltd, Chichester, UK, 2009. https://doi.org/10.1002/9780470743386.

[37] L. V. Hedges, I. Olkin, Statistical methods for meta-analysis, Academic Press, New York, 1985.

[38] S.K. Basu, S. Pradhan, K. Kapse, R. McCarter, J. Murnick, T. Chang, C. Limperopoulos, Third Trimester Cerebellar Metabolite Concentrations are Decreased in Very Premature Infants with Structural Brain Injury, Sci. Rep. 9 (2019). https://doi.org/10.1038/s41598-018-37203-4.

[39] P.E. Sijens, K. Wischniowsky, H.J. ter Horst, The prognostic value of proton magnetic resonance spectroscopy in term newborns treated with therapeutic hypothermia following asphyxia, Magn. Reson. Imaging. 42 (2017) 82–87. https://doi.org/10.1016/j.mri.2017.06.001.

[40] P.J. Lally, P. Montaldo, V. Oliveira, A. Soe, R. Swamy, P. Bassett, J. Mendoza, G. Atreja, U. Kariholu, S. Pattnayak, P. Sashikumar, H. Harizaj, M. Mitchell, V. Ganesh, S. Harigopal, J. Dixon, P. English, P. Clarke, P. Muthukumar, P. Satodia, S. Wayte, L.J. Abernethy, K. Yajamanyam, A. Bainbridge, D. Price, A. Huertas, D.J. Sharp, V. Kalra, S. Chawla, S. Shankaran, S. Thayyil, S. Harigopal, Magnetic resonance spectroscopy assessment of brain injury after moderate hypothermia in neonatal encephalopathy: a prospective multicentre cohort study, Lancet Neurol. 18 (2019) 35–45. https://doi.org/10.1016/S1474-4422(18)30325-9.

[41] R. V. Simões, M. Cruz-Lemini, N. Bargalló, E. Gratacós, M. Sanz-Cortés, Brain metabolite differences in one-year-old infants born small at term and association with neurodevelopmental outcome, Am. J. Obstet. Gynecol. 213 (2015) 210.e1–210.e11. https://doi.org/10.1016/j.ajog.2015.04.011.

[42] F.M. Howells, K.A. Donald, A. Roos, R.P. Woods, H.J. Zar, K.L. Narr, D.J. Stein, Reduced glutamate in white matter of male neonates exposed to alcohol in utero: a 1H-magnetic resonance spectroscopy study, Metab. Brain Dis. 31 (2016) 1105–1112. https://doi.org/10.1007/s11011-016-9850-x.

[43] J. Shibasaki, T. Niwa, A. Piedvache, M. Tomiyasu, N. Morisaki, Y. Fujii, K. Toyoshima, N. Aida, Comparison of Predictive Values of Magnetic Resonance Biomarkers Based on Scan Timing in Neonatal Encephalopathy Following Therapeutic Hypothermia, J. Pediatr. 239 (2021) 101–109.e4. https://doi.org/10.1016/j.jpeds.2021.08.011.

[44] S.K. Basu, S. Pradhan, M.B. Jacobs, M. Said, K. Kapse, J. Murnick, M.T. Whitehead, T. Chang, A.J. du Plessis, C. Limperopoulos, Age and Sex Influences Gamma-aminobutyric Acid Concentrations in the Developing Brain of Very Premature Infants, Sci. Rep. 10 (2020) 10549. https://doi.org/10.1038/s41598-020-67188-y.

[45] D.F. Van Rappard, A. Klauser, M.E. Steenweg, J.J. Boelens, M. Bugiani, M.S. Van Der Knaap, N.I. Wolf, P.J.W. Pouwels, Quantitative MR spectroscopic imaging in metachromatic leukodystrophy: Value for prognosis and treatment, J. Neurol. Neurosurg. Psychiatry. 89 (2018) 105–111. https://doi.org/10.1136/jnnp-2017-316364.

[46] F. Raschke, R. Noeske, R.. Dineen, D.P. Auer, Measuring Cerebral and Cerebellar Glutathione in Children Using 1 H MEGA-PRESS MRS, Am. J. Neuroradiol. 39 (2018) 375–379. https://doi.org/10.3174/ajnr.A5457.

[47] E.M. Mahone, N.A. Puts, R.A.E. Edden, M. Ryan, H.S. Singer, GABA and glutamate in children with Tourette syndrome: A H-1 MR spectroscopy study at 7 T, PSYCHIATRY Res. 273 (2018) 46–53. https://doi.org/10.1016/j.pscychresns.2017.12.005.

[48] M.J. Holmes, F.C. Robertson, F. Little, S.R. Randall, M.F. Cotton, A.J.W. van der Kouwe, B. Laughton, E.M. Meintjes, Longitudinal increases of brain metabolite levels in 5-10 year old children, PLoS One. 12 (2017). https://doi.org/10.1371/journal.pone.0180973.

[49] W.J. Zhang, F.G. Nery, M.J. Tallman, L.R. Patino, C.M. Adler, J.R. Strawn, D.E. Fleck, D.H. Barzman, J.A. Sweeney, S.M. Strakowski, S. Lui, M.P. DelBello, Individual prediction of symptomatic converters in youth offspring of bipolar parents using proton magnetic resonance spectroscopy, Eur. Child Adolesc. Psychiatry. 30 (2021) 55–64. https://doi.org/10.1007/s00787-020-01483-x.

[50] B. Holshouser, J. Pivonka-Jones, J.G. Nichols, U. Oyoyo, K.R. Tong, N. Ghosh, S. Ashwal, Longitudinal Metabolite Changes after Traumatic Brain Injury: A Prospective Pediatric Magnetic Resonance Spectroscopic Imaging Study, J. Neurotrauma. 36 (2019) 1352–1360. https://doi.org/10.1089/neu.2018.5919.

[51] T. Hai, R. Swansburg, C.K. Kahl, H. Frank, J.F. Lemay, F.P. Macmaster, Magnetic Resonance Spectroscopy of γ-Aminobutyric Acid and Glutamate Concentrations in Children with Attention-Deficit/Hyperactivity Disorder, JAMA Netw. Open. 3 (2020) 2–5. https://doi.org/10.1001/jamanetworkopen.2020.20973.

[52] F.G. Nery, W.A. Weber, T.J. Blom, J. Welge, L.R. Patino, J.R. Strawn, W.J. Chu, C.M. Adler, R.A. Komoroski, S.M. Strakowski, M.P. DelBello, Longitudinal proton spectroscopy study of the prefrontal cortex in youth at risk for bipolar disorder before and after their first mood episode, BIPOLAR Disord. 21 (2019) 330–341. https://doi.org/10.1111/bdi.12770.

[53] M. Cichocka, J. Kozub, P. Karcz, A. Urbanik, Sex differences in brain metabolite concentrations in healthy children – proton magnetic resonance spectroscopy study (1 HMRS), Polish J. Radiol. 83 (2018) 24–31. https://doi.org/10.5114/pjr.2018.74536.

[54] L. Calderón-Garcidueñas, A. Mora-Tiscareño, G. Melo-Sánchez, J. Rodríguez-Díaz, R. Torres-Jardón, M. Styner, P.S. Mukherjee, W. Lin, V. Jewells, A critical proton MR spectroscopy marker of Alzheimer’s disease early neurodegenerative change: Low hippocampal NAA/Cr ratio impacts APOE ε 4 Mexico City children and their parents, J. Alzheimer’s Dis. 48 (2015) 1065–1075. https://doi.org/10.3233/JAD-150415.

[55] J. Tannous, B. Cao, J.A. Stanley, G.B. Zunta-Soares, B. Mwangi, M. Sanches, J.C. Soares, Altered neurochemistry in the anterior white matter of bipolar children and adolescents: a multivoxel 1H MRS study, Mol. Psychiatry. 26 (2021) 4117–4126. https://doi.org/10.1038/s41380-020-00927-9.

[56] J.P. Hegarty, M. Gu, D.M. Spielman, S.C. Cleveland, J.F. Hallmayer, L.C. Lazzeroni, M.M. Raman, T.W. Frazier, J.M. Phillips, A.L. Reiss, A.Y. Hardan, A proton MR spectroscopy study of the thalamus in twins with autism spectrum disorder, Prog. Neuro-Psychopharmacology Biol. Psychiatry. 81 (2018) 153–160. https://doi.org/10.1016/j.pnpbp.2017.09.016.

[57] E.T. Wood, K.K. Cummings, J. Jung, G. Patterson, N. Okada, J. Guo, J. O’Neill, M. Dapretto, S.Y. Bookheimer, S.A. Green, Sensory over-responsivity is related to GABAergic inhibition in thalamocortical circuits, Transl. Psychiatry. 11 (2021). https://doi.org/10.1038/s41398-020-01154-0.

[58] W. Wang, H. Sun, X. Su, Q. Tan, S. Zhang, C. Xia, L. Li, G.J. Kemp, Q. Yue, Q. Gong, Increased right amygdala metabolite concentrations in the absence of atrophy in children and adolescents with PTSD, Eur. Child Adolesc. Psychiatry. 28 (2019) 807–817. https://doi.org/10.1007/s00787-018-1241-x.

[59] M.C. Craig, L.M. Mulder, M.P. Zwiers, A. Sethi, P.J. Hoekstra, A. Dietrich, S. Baumeister, P.M. Aggensteiner, T. Banaschewski, D. Brandeis, J.E. Werhahn, S. Walitza, J. Castro-Fornieles, C. Arango, U.M.E. Schulze, J.C. Glennon, B. Franke, P.J. Santosh, M. Mastroianni, J.J.A. van Asten, J.K. Buitelaar, D.J. Lythgoe, J. Naaijen, Distinct associations between fronto-striatal glutamate concentrations and callous-unemotional traits and proactive aggression in disruptive behavior, Cortex. 121 (2019) 135–146. https://doi.org/10.1016/j.cortex.2019.08.017.

[60] T. Horowitz-Kraus, K.J. Brunst, K.M. Cecil, Children With Dyslexia and Typical Readers: Sex-Based Choline Differences Revealed Using Proton Magnetic Resonance Spectroscopy Acquired Within Anterior Cingulate Cortex, Front. Hum. Neurosci. 12 (2018). https://doi.org/10.3389/fnhum.2018.00466.

[61] C. Gasparovic, A. Caprihan, R.A. Yeo, J. Phillips, J.R. Lowe, R. Campbell, R.K. Ohls, The long-term effect of erythropoiesis stimulating agents given to preterm infants: a proton magnetic resonance spectroscopy study on neurometabolites in early childhood, Pediatr. Radiol. 48 (2018) 374–382. https://doi.org/10.1007/s00247-017-4052-1.

[62] H.L. Carlson, F.P. MacMaster, A.D. Harris, A. Kirton, Spectroscopic Biomarkers of Motor Cortex Developmental Plasticity in Hemiparetic Children after Perinatal Stroke, Hum. Brain Mapp. 38 (2017) 1574–1587. https://doi.org/10.1002/hbm.23472.

[63] J. O’Neill, M.J. O’Connor, V. Yee, R. Ly, K. Narr, J.R. Alger, J.G. Levitt, Differential neuroimaging indices in prefrontal white matter in prenatal alcohol-associated ADHD versus idiopathic ADHD, Birth Defects Res. 111 (2019) 797–811. https://doi.org/10.1002/bdr2.1460.

[64] C.A. Steinegger, N. Zoelch, A. Hock, A. Henning, E.J.E. Engeli, E. Seifritz, L.M. Hulka, M. Herdener, Neurometabolic alterations in the nucleus accumbens of smokers assessed with H-1 magnetic resonance spectroscopy: The role of glutamate and neuroinflammation, Addict. Biol. (n.d.). https://doi.org/10.1111/adb.13027.

[65] L.L. Gramegna, A. Pisano, C. Testa, D.N. Manners, R. D’Angelo, E. Boschetti, F. Giancola, L. Pironi, L. Caporali, M. Capristo, M.L. Valentino, G. Plazzi, C. Casali, M.T. Dotti, G. Cenacchi, M. Hirano, C. Giordano, P. Parchi, R. Rinaldi, R. De Giorgio, R. Lodi, V. Carelli, C. Tonon, Cerebral Mitochondrial Microangiopathy Leads to Leukoencephalopathy in Mitochondrial Neurogastrointestinal Encephalopathy., AJNR. Am. J. Neuroradiol. 39 (2018) 427–434. https://doi.org/10.3174/ajnr.A5507.

[66] T.C. Ford, L.A. Downey, T. Simpson, G. McPhee, C. Oliver, C. Stough, The Effect of a High-Dose Vitamin B Multivitamin Supplement on the Relationship between Brain Metabolism and Blood Biomarkers of Oxidative Stress: A Randomized Control Trial, Nutrients. 10 (2018). https://doi.org/10.3390/nu10121860.

[67] J.L. Kroll, A.M. Steele, A.E. Pinkham, C. Choi, D.A. Khan, S. V Patel, J.R. Chen, S. Aslan, E.S. Brown, T. Ritz, Hippocampal metabolites in asthma and their implications for cognitive function, NEUROIMAGE-CLINICAL. 19 (2018) 213–221. https://doi.org/10.1016/j.nicl.2018.04.012.

[68] E. Güleş, D.V. Iosifescu, Ü. Tural, Plasma Neuronal and Glial Markers and Anterior Cingulate Metabolite Levels in Major Depressive Disorder: A Pilot Study, Neuropsychobiology. 79 (2020) 214–221. https://doi.org/10.1159/000505782.

[69] Y. Shan, Y. Jia, S. Zhong, X. Li, H. Zhao, J. Chen, Q. Lu, L. Zhang, Z. Li, S. Lai, Y. Wang, Correlations between working memory impairment and neurometabolites of prefrontal cortex and lenticular nucleus in patients with major depressive disorder, J. Affect. Disord. 227 (2018) 236–242. https://doi.org/10.1016/j.jad.2017.10.030.

[70] O.M. Gonen, B.A. Moffat, P. Kwan, T.J. O’Brien, P.M. Desmond, E. Lui, Reproducibility of Glutamate, Glutathione, and GABA Measurements in vivo by Single-Voxel STEAM Magnetic Resonance Spectroscopy at 7-Tesla in Healthy Individuals, Front. Neurosci. 14 (2020) 1–9. https://doi.org/10.3389/fnins.2020.566643.

[71] S. Singh, S. Khushu, P. Kumar, S. Goyal, T. Bhatia, S.N. Deshpande, Evidence for regional hippocampal damage in patients with schizophrenia, Neuroradiology. 60 (2018) 199–205. https://doi.org/10.1007/s00234-017-1954-4.

[72] M. Dehghani, K.Q. Do, P. Magistretti, L. Xin, Lactate measurement by neurochemical profiling in the dorsolateral prefrontal cortex at 7T: accuracy, precision, and relaxation times, Magn. Reson. Med. 83 (2020) 1895–1908. https://doi.org/10.1002/mrm.28066.

[73] S. Chawla, S.C. Lee, S. Mohan, S. Wang, M. Nasrallah, A. Vossough, J. Krejza, E.R. Melhem, S. Ali Nabavizadeh, Lack of choline elevation on proton magnetic resonance spectroscopy in grade I–III gliomas, Neuroradiol. J. 32 (2019) 250–258. https://doi.org/10.1177/1971400919846509.

[74] G.H. Kim, I. Kang, H. Jeong, S. Park, H. Hong, J. Kim, J.Y. Kim, R.A.E. Edden, I.K. Lyoo, S. Yoon, Low Prefrontal GABA Levels Are Associated With Poor Cognitive Functions in Professional Boxers, Front. Hum. Neurosci. 13 (2019). https://doi.org/10.3389/fnhum.2019.00193.

[75] H. Su, T. Chen, N. Zhong, H. Jiang, J. Du, K. Xiao, D. Xu, W. Song, M. Zhao, Decreased GABA concentrations in left prefrontal cortex of methamphetamine dependent patients: A proton magnetic resonance spectroscopy study, J. Clin. Neurosci. 71 (2020) 15–20. https://doi.org/10.1016/j.jocn.2019.11.021.

[76] T.L. White, M.A. Monnig, E.G. Walsh, A.Z. Nitenson, A.D. Harris, R.A. Cohen, E.C. Porges, A.J. Woods, D.G. Lamb, C.A. Boyd, S. Fekir, Psychostimulant drug effects on glutamate, Glx, and creatine in the anterior cingulate cortex and subjective response in healthy humans, NEUROPSYCHOPHARMACOLOGY. 43 (2018) 1498–1509. https://doi.org/10.1038/s41386-018-0027-7.

[77] J.T. Kantrowitz, Z. Dong, M.S. Milak, R. Rashid, L.S. Kegeles, D.C. Javitt, J.A. Lieberman, J. John Mann, Ventromedial prefrontal cortex/anterior cingulate cortex Glx, glutamate, and GABA levels in medication-free major depressive disorder, Transl. Psychiatry. 11 (2021) 1–6. https://doi.org/10.1038/s41398-021-01541-1.

[78] C.M. Moon, G.W. Jeong, Associations of neurofunctional, morphometric and metabolic abnormalities with clinical symptom severity and recognition deficit in obsessive–compulsive disorder, J. Affect. Disord. 227 (2018) 603–612. https://doi.org/10.1016/j.jad.2017.11.059.

[79] N.W. Duncan, J. Zhang, G. Northoff, X. Weng, Investigating GABA concentrations measured with macromolecule suppressed and unsuppressed MEGA-PRESS MR spectroscopy and their relationship with BOLD responses in the occipital cortex, J. Magn. Reson. Imaging. 50 (2019) 1285–1294. https://doi.org/10.1002/jmri.26706.

[80] M.G. Bossong, M. Antoniades, M. Azis, C. Samson, B. Quinn, I. Bonoldi, G. Modinos, J. Perez, O.D. Howes, J.M. Stone, P. Allen, P. McGuire, Association of Hippocampal Glutamate Levels with Adverse Outcomes in Individuals at Clinical High Risk for Psychosis, JAMA Psychiatry. 76 (2019) 199–207. https://doi.org/10.1001/jamapsychiatry.2018.3252.

[81] B. Schmitz, X. Wang, P.B. Barker, U. Pilatus, P. Bronzlik, M. Dadak, K.G. Kahl, H. Lanfermann, X.Q. Ding, Effects of Aging on the Human Brain: A Proton and Phosphorus MR Spectroscopy Study at 3T, J. NEUROIMAGING. 28 (2018) 416–421. https://doi.org/10.1111/jon.12514.

[82] J.M. Coughlin, K. Yang, A. Marsman, S. Pradhan, M. Wang, R.E. Ward, S. Bonekamp, E.B. Ambinder, C.P. Higgs, P.K. Kim, J.A. Edwards, M. Varvaris, H. Wang, S. Posporelis, S. Ma, T. Tsujimura, R.A.E. Edden, M.G. Pomper, T.W. Sedlak, M. Fournier, D.J. Schretlen, N.G. Cascella, P.B. Barker, A. Sawa, A multimodal approach to studying the relationship between peripheral glutathione, brain glutamate, and cognition in health and in schizophrenia, Mol. Psychiatry. 26 (2021) 3502–3511. https://doi.org/10.1038/s41380-020-00901-5.

[83] I.A. Giapitzakis, T.T. Shao, N. Avdievich, R. Mekle, R. Kreis, A. Henning, Metabolite-cycled STEAM and semi-LASER localization for MR spectroscopy of the human brain at 9.4T, Magn. Reson. Med. 79 (2018) 1841–1850. https://doi.org/10.1002/mrm.26873.

[84] C.M. Kaplan, A. Schrepf, D. Vatansever, T.E. Larkin, I. Mawla, E. Ichesco, L. Kochlefl, S.E. Harte, D.J. Clauw, G.A. Mashour, R.E. Harris, Functional and neurochemical disruptions of brain hub topology in chronic pain, Pain. 160 (2019) 973–983. https://doi.org/10.1097/j.pain.0000000000001480.

[85] L. An, M.F. Araneta, C. Johnson, J. Shen, Simultaneous measurement of glutamate, glutamine, GABA, and glutathione by spectral editing without subtraction, Magn. Reson. Med. 80 (2018) 1776–1786. https://doi.org/10.1002/mrm.27172.

[86] J. Lynn, E.A. Woodcock, C. Anand, D. Khatib, J.A. Stanley, Differences in steady-state glutamate levels and variability between “non-task-active” conditions: Evidence from H-1 fMRS of the prefrontal cortex, Neuroimage. 172 (2018) 554–561. https://doi.org/10.1016/j.neuroimage.2018.01.069.

[87] T.P. Lawrence, A. Steel, M. Ezra, M. Speirs, P.M. Pretorius, G. Douaud, S. Sotiropoulos, T. Cadoux-Hudson, U.E. Emir, N.L. Voets, MRS and DTI evidence of progressive posterior cingulate cortex and corpus callosum injury in the hyper-acute phase after Traumatic Brain Injury, BRAIN Inj. 33 (2019) 854–868. https://doi.org/10.1080/02699052.2019.1584332.

[88] F. Caravaggio, Y. Iwata, E. Plitman, S. Chavez, C. Borlido, J.K. Chung, J. Kim, M. Agarwal, P. Gerretsen, G. Remington, M. Hahn, A. Graff-Guerrero, Reduced insulin sensitivity may be related to less striatal glutamate: An H-1-MRS study in healthy non-obese humans, Eur. Neuropsychopharmacol. 28 (2018) 285–296. https://doi.org/10.1016/j.euroneuro.2017.12.002.

[89] T.L. White, M.A. Gonsalves, R.A. Cohen, A.D. Harris, M.A. Monnig, E.G. Walsh, A.Z. Nitenson, E.C. Porges, D.G. Lamb, A.J. Woods, C.B. Borja, The neurobiology of wellness: H-1-MRS correlates of agency, flexibility and neuroaffective reserves in healthy young adults, Neuroimage. 225 (2021). https://doi.org/10.1016/j.neuroimage.2020.117509.

[90] C. Volk, V. Jaramillo, R. Merki, R.O. Tuura, R. Huber, Diurnal changes in glutamate plus glutamine levels of healthy young adults assessed by proton magnetic resonance spectroscopy, Hum. Brain Mapp. 39 (2018) 3984–3992. https://doi.org/10.1002/hbm.24225.

[91] G. Cao, R.A.E. Edden, F. Gao, H. Li, T. Gong, W. Chen, X. Liu, G. Wang, B. Zhao, Reduced GABA levels correlate with cognitive impairment in patients with relapsing-remitting multiple sclerosis, Eur. Radiol. 28 (2018) 1140–1148. https://doi.org/10.1007/s00330-017-5064-9.

[92] H. Cen, J. Xu, Z. Yang, L. Mei, T. Chen, K. Zhuo, Q. Xiang, Z. Song, Y. Wang, X. Guo, J. Wang, K. Jiang, Y. Xu, Y. Li, D. Liu, Neurochemical and brain functional changes in the ventromedial prefrontal cortex of first-episode psychosis patients: A combined functional magnetic resonance imaging—proton magnetic resonance spectroscopy study, Aust. N. Z. J. Psychiatry. 54 (2020) 519–527. https://doi.org/10.1177/0004867419898520.

[93] H.J. Patel, S. Romanzetti, A. Pellicano, M.A. Nitsche, K. Reetz, F. Binkofski, Proton Magnetic Resonance Spectroscopy of the motor cortex reveals long term GABA change following anodal Transcranial Direct Current Stimulation, Sci. Rep. 9 (2019). https://doi.org/10.1038/s41598-019-39262-7.

[94] J. Kumar, E.B. Liddle, C.C. Fernandes, L. Palaniyappan, E.L. Hall, S.E. Robson, M. Simmonite, J. Fiesal, M.Z. Katshu, A. Qureshi, M. Skelton, N.G. Christodoulou, M.J. Brookes, P.G. Morris, P.F. Liddle, Glutathione and glutamate in schizophrenia: a 7T MRS study, Mol. Psychiatry. 25 (2020) 873–882. https://doi.org/10.1038/s41380-018-0104-7.

[95] M. Corcoran, E.L. Hawkins, D. O’Hora, H.C. Whalley, J. Hall, S.M. Lawrie, M.R. Dauvermann, Are working memory and glutamate concentrations involved in early-life stress and severity of psychosis?, Brain Behav. 10 (2020) 1–13. https://doi.org/10.1002/brb3.1616.

[96] A. Burger, S.J. Brooks, D.J. Stein, F.M. Howells, The impact of acute and short-term methamphetamine abstinence on brain metabolites: A proton magnetic resonance spectroscopy chemical shift imaging study, Drug Alcohol Depend. 185 (2018) 226–237. https://doi.org/10.1016/j.drugalcdep.2017.11.029.

[97] S. Suri, U. Emir, C.J. Stagg, J. Near, R. Mekle, F. Schubert, E. Zsoldos, A. Mahmood, A. Singh-Manoux, M. Kivimaki, K.P. Ebmeier, C.E. Mackay, N. Filippini, Effect of age and the APOE gene on metabolite concentrations in the posterior cingulate cortex, Neuroimage. 152 (2017) 509–516. https://doi.org/10.1016/j.neuroimage.2017.03.031.

[98] A.L. Yasen, M.M. Lim, K.B. Weymann, A.D. Christie, Excitability, Inhibition, and Neurotransmitter Levels in the Motor Cortex of Symptomatic and Asymptomatic Individuals Following Mild Traumatic Brain Injury, Front. Neurol. 11 (2020). https://doi.org/10.3389/fneur.2020.00683.

[99] X.Q. Ding, A.A. Maudsley, U. Schweiger, B. Schmitz, R. Lichtinghagen, S. Bleich, H. Lanfermann, K.G. Kahl, Effects of a 72 hours fasting on brain metabolism in healthy women studied in vivo with magnetic resonance spectroscopic imaging, J. Cereb. BLOOD FLOW Metab. 38 (2018) 469–478. https://doi.org/10.1177/0271678X17697721.

[100] M.İ. Atagün, E.M. Şıkoğlu, Ç. Soykan, C. Serdar Süleyman, S. Ulusoy-Kaymak, A. Çayköylü, O. Algın, M.L. Phillips, D. Öngür, C.M. Moore, Perisylvian GABA levels in schizophrenia and bipolar disorder, Neurosci. Lett. 637 (2017) 70–74. https://doi.org/10.1016/j.neulet.2016.11.051.

[101] A.L. Yasen, G.N. Eick, K.N. Sterner, A.D. Christie, Motor Cortex Function in APOE4 Carriers and Noncarriers, J. Clin. Neurophysiol. 38 (2021) 553–557. https://doi.org/10.1097/WNP.0000000000000738.

[102] A. Strasser, L.J. Xin, R. Gruetter, C. Sandi, Nucleus accumbens neurochemistry in human anxiety: A 7 T H-1-MRS study, Eur. Neuropsychopharmacol. 29 (2019) 365–375. https://doi.org/10.1016/j.euroneuro.2018.12.015.

[103] M.S. Davitz, O. Gonen, A. Tal, J.S. Babb, Y.W. Lui, I.I. Kirov, Quantitative multivoxel proton MR spectroscopy for the identification of white matter abnormalities in mild traumatic brain injury: Comparison between regional and global analysis, J. Magn. Reson. IMAGING. 50 (2019) 1424–1432. https://doi.org/10.1002/jmri.26718.

[104] J. Horder, M.M. Petrinovic, M.A. Mendez, A. Bruns, T. Takumi, W. Spooren, G.J. Barker, B. Kunnecke, D.G. Murphy, Glutamate and GABA in autism spectrum disorder-a translational magnetic resonance spectroscopy study in man and rodent models, Transl. Psychiatry. 8 (2018). https://doi.org/10.1038/s41398-018-0155-1.

[105] A.A. Bhogal, T.A.A. Broeders, L. Morsinkhof, M. Edens, S. Nassirpour, P. Chang, D.W.J. Klomp, C.H. Vinkers, J.P. Wijnen, Lipid-suppressed and tissue-fraction corrected metabolic distributions in human central brain structures using 2D H-1 magnetic resonance spectroscopic imaging at 7 T, BRAIN Behav. 10 (2020). https://doi.org/10.1002/brb3.1852.

[106] H. Hjelmervik, A.R. Craven, I. Sinceviciute, E. Johnsen, K. Kompus, J.J. Bless, R.A. Kroken, E.M. Løberg, L. Ersland, R. Grüner, K. Hugdahl, Intra-Regional Glu-GABA vs Inter-Regional Glu-Glu Imbalance: A 1H-MRS Study of the Neurochemistry of Auditory Verbal Hallucinations in Schizophrenia, Schizophr. Bull. 46 (2020) 633–642. https://doi.org/10.1093/schbul/sbz099.

[107] M.G. Soeiro-de-Souza, E. Scotti-Muzzi, F. Fernandes, R.T. De Sousa, C.C. Leite, M.C. Otaduy, R. Machado-Vieira, Anterior cingulate cortex neuro-metabolic changes underlying lithium-induced euthymia in bipolar depression: A longitudinal H-1-MRS study, Eur. Neuropsychopharmacol. 49 (2021) 93–100. https://doi.org/10.1016/j.euroneuro.2021.03.020.

[108] J. Wang, T. Zhou, J. Liu, J. Shangguan, X. Liu, Z. Li, X. Zhou, Y. Ren, C. Wang, Application of 1H-MRS in end-stage renal disease with depression, BMC Nephrol. 21 (2020) 225. https://doi.org/10.1186/s12882-020-01863-0.

[109] M. Rogdaki, P. Hathway, M. Gudbrandsen, R.A. McCutcheon, S. Jauhar, E. Daly, O. Howes, Glutamatergic function in a genetic high-risk group for psychosis: A proton magnetic resonance spectroscopy study in individuals with 22q11.2 deletion, Eur. Neuropsychopharmacol. 29 (2019) 1333–1342. https://doi.org/10.1016/j.euroneuro.2019.09.005.

[110] T.G. Stærmose, M.K. Knudsen, H. Kasch, J.U. Blicher, Cortical GABA in migraine with aura - an ultrashort echo magnetic resonance spectroscopy study, J. Headache Pain. 20 (2019). https://doi.org/10.1186/s10194-019-1059-z.

[111] M. Xia, J. Wang, J. Sheng, Y. Tang, C. Li, K. Lim, B. He, C. Li, Y. Xu, J. Wang, Effect of Electroconvulsive Therapy on Medial Prefrontal γ-Aminobutyric Acid among Schizophrenia Patients: A Proton Magnetic Resonance Spectroscopy Study, J. ECT. 34 (2018) 227–232. https://doi.org/10.1097/YCT.0000000000000507.

[112] J.J. Prisciandaro, M. Mikkelsen, M.G. Saleh, R.A.E. Edden, An evaluation of the reproducibility of H-1-MRS GABA and GSH levels acquired in healthy volunteers with J-difference editing sequences at varying echo times, Magn. Reson. Imaging. 65 (2020) 109–113. https://doi.org/10.1016/j.mri.2019.10.004.

[113] Y.M. Chan, K. Pitchaimuthu, Q.Z. Wu, O.L. Carter, G.F. Egan, D.R. Badcock, A.M. McKendrick, Relating excitatory and inhibitory neurochemicals to visual perception: A magnetic resonance study of occipital cortex between migraine events, PLoS One. 14 (2019) 1–13. https://doi.org/10.1371/journal.pone.0208666.

[114] R. Zawadzki, B. Kubas, M. Hładuński, O. Zajkowska, J. Zajkowska, D. Jurgilewicz, A. Garkowski, S. Pancewicz, U. Łebkowska, Proton magnetic resonance spectroscopy (1 H-MRS) of the brain in patients with tick-borne encephalitis, Sci. Rep. 9 (2019) 1–6. https://doi.org/10.1038/s41598-019-39352-6.

[115] M. Považan, B. Strasser, G. Hangel, E. Heckova, S. Gruber, S. Trattnig, W. Bogner, Simultaneous mapping of metabolites and individual macromolecular components via ultra-short acquisition delay 1H MRSI in the brain at 7T, Magn. Reson. Med. 79 (2018) 1231–1240. https://doi.org/10.1002/mrm.26778.

[116] R.S. Huber, D.G. Kondo, X.F. Shi, A.P. Prescot, E. Clark, P.F. Renshaw, D.A. Yurgelun-Todd, Relationship of executive functioning deficits to N-acetyl aspartate (NAA) and gamma-aminobutyric acid (GABA) in youth with bipolar disorder, J. Affect. Disord. 225 (2018) 71–78. https://doi.org/10.1016/j.jad.2017.07.052.

[117] J. Bauer, A. Werner, W. Kohl, H. Kugel, A. Shushakova, A. Pedersen, P. Ohrmann, Hyperactivity and impulsivity in adult attention-deficit/hyperactivity disorder is related to glutamatergic dysfunction in the anterior cingulate cortex, World J. Biol. Psychiatry. 19 (2018) 538–546. https://doi.org/10.1080/15622975.2016.1262060.

[118] M.D. Legarreta, C. Sheth, A.P. Prescot, P.F. Renshaw, E.C. McGlade, D.A. Yurgelun-Todd, An exploratory proton MRS examination of gamma-aminobutyric acid, glutamate, and glutamine and their relationship to affective aspects of chronic pain, Neurosci. Res. 163 (2021) 10–17. https://doi.org/10.1016/j.neures.2020.03.002.

[119] R. Wang, B. Hu, C. Sun, D. Geng, J. Lin, Y. Li, Metabolic abnormality in acute stroke-like lesion and its relationship with focal cerebral blood flow in patients with MELAS: Evidence from proton MR spectroscopy and arterial spin labeling, Mitochondrion. 59 (2021) 276–282. https://doi.org/10.1016/j.mito.2021.06.012.

[120] A. Parmar, P. Sharan, S.K. Khandelwal, K. Agarwal, U. Sharma, N.R. Jagannathan, Brain neurochemistry in unmedicated obsessive–compulsive disorder patients and effects of 12-week escitalopram treatment: 1H-magnetic resonance spectroscopy study, Psychiatry Clin. Neurosci. 73 (2019) 386–393. https://doi.org/10.1111/pcn.12850.

[121] I.M. Adanyeguh, M.L. Monin, D. Rinaldi, L. Freeman, A. Durr, S. Lehericy, P.G. Henry, F. Mochel, Expanded neurochemical profile in the early stage of Huntington disease using proton magnetic resonance spectroscopy, NMR Biomed. 31 (2018). https://doi.org/10.1002/nbm.3880.

[122] M. Gajdošík, K. Landheer, K.M. Swanberg, F. Adlparvar, G. Madelin, W. Bogner, C. Juchem, I.I. Kirov, Hippocampal single-voxel MR spectroscopy with a long echo time at 3 T using semi-LASER sequence, NMR Biomed. 34 (2021) 1–14. https://doi.org/10.1002/nbm.4538.

[123] Y. Koush, R.A. de Graaf, L. Jiang, D.L. Rothman, F. Hyder, Functional MRS with J-edited lactate in human motor cortex at 4 T, Neuroimage. 184 (2019) 101–108. https://doi.org/10.1016/j.neuroimage.2018.09.008.

[124] M. Draganov, Y. Vives-Gilabert, J. de Diego-Adeliño, M. Vicent-Gil, D. Puigdemont, M.J. Portella, Glutamatergic and GABA-ergic abnormalities in First-episode depression. A 1-year follow-up 1H-MR spectroscopic study, J. Affect. Disord. 266 (2020) 572–577. https://doi.org/10.1016/j.jad.2020.01.138.

[125] O.E. Rowe, D. Rangaprakash, A. Weerasekera, N. Godbole, E. Haxton, P.F. James, C.D. Stephen, R.L. Barry, F.S. Eichler, E.M. Ratai, Magnetic resonance imaging and spectroscopy in late-onset GM2-gangliosidosis, Mol. Genet. Metab. 133 (2021) 386–396. https://doi.org/10.1016/j.ymgme.2021.06.008.

[126] H. Hjelmervik, M. Hausmann, A.R. Craven, M. Hirnstein, K. Hugdahl, K. Specht, Sex- and sex hormone-related variations in energy-metabolic frontal brain asymmetries: A magnetic resonance spectroscopy study, Neuroimage. 172 (2018) 817–825. https://doi.org/10.1016/j.neuroimage.2018.01.043.

[127] H.J. Zöllner, M. Považan, S.C.N. Hui, S. Tapper, R.A.E. Edden, G. Oeltzschner, Comparison of different linear-combination modeling algorithms for short-TE proton spectra, NMR Biomed. 34 (2021) 1–17. https://doi.org/10.1002/nbm.4482.

[128] E.J. Cadena, D.M. White, N. V Kraguljac, M.A. Reid, J.O. Maximo, E.A. Nelson, B.A. Gawronski, A.C. Lahti, A Longitudinal Multimodal Neuroimaging Study to Examine Relationships Between Resting State Glutamate and Task Related BOLD Response in Schizophrenia, Front. PSYCHIATRY. 9 (2018). https://doi.org/10.3389/fpsyt.2018.00632.

[129] K.G. Kahl, S. Atalay, A.A. Maudsley, S. Sheriff, A. Cummings, H. Frieling, B. Schmitz, H. Lanfermann, X.Q. Ding, Altered neurometabolism in major depressive disorder: A whole brain 1H-magnetic resonance spectroscopic imaging study at 3T, Prog. Neuro-Psychopharmacology Biol. Psychiatry. 101 (2020). https://doi.org/10.1016/j.pnpbp.2020.109916.

[130] M.G. Bossong, R. Wilson, E. Appiah-Kusi, P. McGuire, S. Bhattacharyya, Human Striatal Response to Reward Anticipation Linked to Hippocampal Glutamate Levels, Int. J. Neuropsychopharmacol. 21 (2018) 623–630. https://doi.org/10.1093/ijnp/pyy011.

[131] P.E. Menshchikov, N.A. Semenova, A. V Manzhurtsev, T.A. Akhadov, S.D. Varfolomeev, Cerebral quantification of N-acetyl aspartate, aspartate, and glutamate levels in local structures of the human brain using J-editing of H-1 magnetic resonance spectra in vivo, Russ. Chem. Bull. 67 (2018) 655–662. https://doi.org/10.1007/s11172-018-2119-2.

[132] Y.Y. Shih, M. Büchert, H.W. Chung, J. Hennig, D. Von Elverfeldt, Vitamin C estimation with standard 1H spectroscopy using a clinical 3T MR system: Detectability and reliability within the human brain, J. Magn. Reson. Imaging. 28 (2008) 351–358. https://doi.org/10.1002/jmri.21466.

[133] P. Bednarik, B. Spurny, L.R. Silberbauer, A. Svatkova, P.A. Handschuh, B. Reiter, M.E. Konadu, T. Stimpfl, M. Spies, W. Bogner, R. Lanzenberger, Effect of Ketamine on Human Neurochemistry in Posterior Cingulate Cortex: A Pilot Magnetic Resonance Spectroscopy Study at 3 Tesla, Front. Neurosci. 15 (2021). https://doi.org/10.3389/fnins.2021.609485.

[134] M.A.P. Bloomfield, K. Petrilli, R. Lees, C. Hindocha, K. Beck, R.J. Turner, E.C. Onwordi, N. Rane, D.J. Lythgoe, J.M. Stone, H.V. Curran, O.D. Howes, T.P. Freeman, The Effects of Acute Δ9-Tetrahydrocannabinol on Striatal Glutamatergic Function: A Proton Magnetic Resonance Spectroscopy Study, Biol. Psychiatry Cogn. Neurosci. Neuroimaging. 6 (2021) 660–667. https://doi.org/10.1016/j.bpsc.2021.04.013.

[135] C. Vingerhoets, D.H.Y. Tse, M. van Oudenaren, D. Hernaus, E. van Duin, J. Zinkstok, J.G. Ramaekers, J.F.A. Jansen, G. McAlonan, T. van Amelsvoort, Glutamatergic and GABAergic reactivity and cognition in 22q11.2 deletion syndrome and healthy volunteers: A randomized double-blind 7-Tesla pharmacological MRS study, J. Psychopharmacol. 34 (2020) 856–863. https://doi.org/10.1177/0269881120922977.

[136] G. Modinos, F. Simsek, J. Horder, M. Bossong, I. Bonoldi, M. Azis, J. Perez, M. Broome, D.J. Lythgoe, J.M. Stone, O.D. Howes, D.G. Murphy, A.A. Grace, P. Allen, P. McGuire, Cortical GABA in subjects at ultra-high risk of psychosis: Relationship to negative prodromal symptoms, Int. J. Neuropsychopharmacol. 21 (2018) 114–119. https://doi.org/10.1093/ijnp/pyx076.

[137] G. Blest-Hopley, A. O’Neill, R. Wilson, V. Giampietro, D. Lythgoe, A. Egerton, S. Bhattacharyya, Adolescent-onset heavy cannabis use associated with significantly reduced glial but not neuronal markers and glutamate levels in the hippocampus, Addict. Biol. 25 (2020). https://doi.org/10.1111/adb.12827.

[138] K.M. Swanberg, H. Prinsen, K. DeStefano, M. Bailey, A. V Kurada, D. Pitt, R.K. Fulbright, C. Juchem, In vivo evidence of differential frontal cortex metabolic abnormalities in progressive and relapsing-remitting multiple sclerosis, NMR Biomed. (n.d.). https://doi.org/10.1002/nbm.4590.

[139] Ş. İ, K.O. N, S.V. G, Study on Dorsolateral Prefrontal Cortex Neurochemical Metabolite Levels of Patients with Major Depression Using H-MRS Technique., Turk Psikiyatri Derg. 31 (2020) 75–83. https://pubmed.ncbi.nlm.nih.gov/32594494/.

[140] J. Kaminski, T. Gleich, Y. Fukuda, T. Katthagen, J. Gallinat, A. Heinz, F. Schlagenhauf, Association of Cortical Glutamate and Working Memory Activation in Patients With Schizophrenia: A Multimodal Proton Magnetic Resonance Spectroscopy and Functional Magnetic Resonance Imaging Study, Biol. Psychiatry. 87 (2020) 225–233. https://doi.org/10.1016/j.biopsych.2019.07.011.

[141] X. Su, C. Xia, W. Wang, H. Sun, Q. Tan, S. Zhang, L. Li, G.J. Kemp, Q. Yue, Q. Gong, Abnormal metabolite concentrations and amygdala volume in patients with recent-onset posttraumatic stress disorder, J. Affect. Disord. 241 (2018) 539–545. https://doi.org/10.1016/j.jad.2018.08.018.

[142] Z. An, V. Tiwari, S.K. Ganji, J. Baxter, M. Levy, M.C. Pinho, E. Pan, E.A. Maher, T.R. Patel, B.E. Mickey, C. Choi, Echo-planar spectroscopic imaging with dual-readout alternated gradients (DRAG-EPSI) at 7 T: Application for 2-hydroxyglutarate imaging in glioma patients, Magn. Reson. Med. 79 (2018) 1851–1861. https://doi.org/10.1002/mrm.26884.

[143] A.G. Costigan, K. Umla-Runge, C.J. Evans, C.J. Hodgetts, A.D. Lawrence, K.S. Graham, Neurochemical correlates of scene processing in the precuneus/posterior cingulate cortex: A multimodal fMRI and 1H-MRS study, Hum. Brain Mapp. 40 (2019) 2884–2898. https://doi.org/10.1002/hbm.24566.

[144] A. Smaragdi, S. Chavez, N.J. Lobaugh, J.H. Meyer, N.J. Kolla, Differential levels of prefrontal cortex glutamate plus glutamine in adults with antisocial personality disorder and bipolar disorder: A proton magnetic resonance spectroscopy study, Prog. Neuropsychopharmacol. Biol. Psychiatry. 93 (2019) 250–255. https://doi.org/10.1016/j.pnpbp.2019.04.002.

[145] H. Maeshima, C. Hosoda, K. Okanoya, T. Nakai, Reduced ?-aminobutyric acid in the superior temporal gyrus is associated with absolute pitch, Neuroreport. 29 (2018) 1487–1491. https://doi.org/10.1097/WNR.0000000000001137.

[146] S. Shakory, J.J. Watts, S. Hafizi, T. Da Silva, S. Khan, M. Kiang, R.M. Bagby, S. Chavez, R. Mizrahi, Hippocampal glutamate metabolites and glial activation in clinical high risk and first episode psychosis, NEUROPSYCHOPHARMACOLOGY. 43 (2018) 2249–2255. https://doi.org/10.1038/s41386-018-0163-0.

[147] J. Archibald, E.L. MacMillan, C. Graf, P. Kozlowski, C. Laule, J.L.K. Kramer, Metabolite activity in the anterior cingulate cortex during a painful stimulus using functional MRS, Sci. Rep. 10 (2020). https://doi.org/10.1038/s41598-020-76263-3.

[148] I.I. Kirov, R. Kuzniecky, H.P. Hetherington, B.J. Soher, M.S. Davitz, J.S. Babb, H.R. Pardoe, J.W. Pan, O. Gonen, Whole brain neuronal abnormalities in focal quantified with proton MR spectroscopy, EPILEPSY Res. 139 (2018) 85–91. https://doi.org/10.1016/j.eplepsyres.2017.11.017.

[149] H. Polacek, E. Kantorova, P. Hnilicova, M. Grendar, K. Zelenak, E. Kurca, Increased glutamate and deep brain atrophy can predict the severity of multiple sclerosis, Biomed. Pap. 163 (2019) 45–53. https://doi.org/10.5507/bp.2018.036.

[150] B. Galińska-Skok, A. Małus, B. Konarzewska, A. Rogowska-Zach, R. Milewski, E. Tarasów, A. Szulc, N. Waszkiewicz, Choline compounds of the frontal lobe and temporal glutamatergic system in bipolar and schizophrenia proton magnetic resonance spectroscopy study, Dis. Markers. 2018 (2018) 1–8. https://doi.org/10.1155/2018/3654894.

[151] M. Colizzi, N. Weltens, P. McGuire, D. Lythgoe, S. Williams, L. Van Oudenhove, S. Bhattacharyya, Delta-9-tetrahydrocannabinol increases striatal glutamate levels in healthy individuals: implications for psychosis, Mol. Psychiatry. 25 (2020) 3231–3240. https://doi.org/10.1038/s41380-019-0374-8.

[152] A. Lind, C.J. Boraxbekk, E.T. Petersen, O.B. Paulson, O. Andersen, H.R. Siebner, A. Marsman, Do glia provide the link between low-grade systemic inflammation and normal cognitive ageing? A H-1 magnetic resonance spectroscopy study at 7 tesla, J. Neurochem. (n.d.). https://doi.org/10.1111/jnc.15456.

[153] E.O. Eroğlu, A. Ertuğrul, K.K. Oğuz, S. Karahan, M.K. Yazici, Effect of Clozapine on Proton Magnetic Resonance Spectroscopy Findings in Hippocampus, Turk Psikiyatr. Derg. 31 (2020) 157–169. https://doi.org/10.5080/u25195.

[154] S. Poletti, M.G. Mazza, B. Vai, C. Lorenzi, C. Colombo, F. Benedetti, Proinflammatory Cytokines Predict Brain Metabolite Concentrations in the Anterior Cingulate Cortex of Patients With Bipolar Disorder, Front. Psychiatry. 11 (2020) 1–9. https://doi.org/10.3389/fpsyt.2020.590095.

[155] M.A. Reid, N. Salibi, D.M. White, T.J. Gawne, T.S. Denney, A.C. Lahti, 7T Proton Magnetic Resonance Spectroscopy of the Anterior Cingulate Cortex in First-Episode Schizophrenia, Schizophr. Bull. 45 (2019) 180–189. https://doi.org/10.1093/schbul/sbx190.

[156] O. Levin, A. Weerasekera, B.R. King, K.F. Heise, D.M. Sima, S. Chalavi, C. Maes, R. Peeters, S. Sunaert, K. Cuypers, S. Van Huffel, D. Mantini, U. Himmelreich, S.P. Swinnen, Sensorimotor cortex neurometabolite levels as correlate of motor performance in normal aging: evidence from a H-1-MRS study, Neuroimage. 202 (2019). https://doi.org/10.1016/j.neuroimage.2019.116050.

[157] C. Wyss, D.H.Y. Tse, F. Boers, N.J. Shah, I. Neuner, W. Kawohl, Association between Cortical GABA and Loudness Dependence of Auditory Evoked Potentials (LDAEP) in Humans, Int. J. Neuropsychopharmacol. 21 (2018) 809–813. https://doi.org/10.1093/ijnp/pyy056.

[158] A. Manzhurtsev, P. Menschchikov, A. Yakovlev, M. Ublinskiy, O. Bozhko, D. Kupriyanov, T. Akhadov, S. Varfolomeev, N. Semenova, 3T MEGA-PRESS study of N-acetyl aspartyl glutamate and N-acetyl aspartate in activated visual cortex, Magn. Reson. Mater. Physics, Biol. Med. 34 (2021) 555–568. https://doi.org/10.1007/s10334-021-00912-5.

[159] M. Marjańska, J.R. McCarten, J. Hodges, L.S. Hemmy, A. Grant, D.K. Deelchand, M. Terpstra, Region-specific aging of the human brain as evidenced by neurochemical profiles measured noninvasively in the posterior cingulate cortex and the occipital lobe using 1H magnetic resonance spectroscopy at 7 T, Neuroscience. 354 (2017) 168–177. https://doi.org/10.1016/j.neuroscience.2017.04.035.

[160] B.R. Godlewska, C. Masaki, A.L. Sharpley, P.J. Cowen, U.E. Emir, Brain glutamate in medication-free depressed patients: a proton MRS study at 7 Tesla, Psychol. Med. 48 (2018) 1731–1737. https://doi.org/10.1017/S0033291717003373.

[161] V. Veeramuthu, P. Seow, V. Narayanan, J.H.D. Wong, L.K. Tan, A.T. Hernowo, N. Ramli, Neurometabolites Alteration in the Acute Phase of Mild Traumatic Brain Injury (mTBI): An in Vivo Proton Magnetic Resonance Spectroscopy (1H-MRS) Study, Acad. Radiol. 25 (2018) 1167–1177. https://doi.org/10.1016/j.acra.2018.01.005.

[162] S. Posporelis, J.M. Coughlin, A. Marsman, S. Pradhan, T. Tanaka, H. Wang, M. Varvaris, R. Ward, C. Higgs, J.A. Edwards, C.N. Ford, P.K. Kim, A.M. Lloyd, R.A.E. Edden, D.J. Schretlen, N.G. Cascella, P.B. Barker, A. Sawa, Decoupling of Brain Temperature and Glutamate in Recent Onset of Schizophrenia: A 7T Proton Magnetic Resonance Spectroscopy Study, Biol. Psychiatry Cogn. Neurosci. Neuroimaging. 3 (2018) 248–254. https://doi.org/10.1016/j.bpsc.2017.04.003.

[163] S.A. Wijtenburg, J. West, S.A. Korenic, F. Kuhney, F.E. Gaston, H.J. Chen, L.M. Rowland, Multimodal Neuroimaging Study of Visual Plasticity in Schizophrenia, Front. PSYCHIATRY. 12 (2021). https://doi.org/10.3389/fpsyt.2021.644271.

[164] C. Wenneberg, M. Nordentoft, E. Rostrup, L.B. Glenthøj, K.B. Bojesen, B. Fagerlund, C. Hjorthøj, K. Krakauer, T.D. Kristensen, C. Schwartz, R.A.E. Edden, B.V. Broberg, B.Y. Glenthøj, Cerebral Glutamate and Gamma-Aminobutyric Acid Levels in Individuals at Ultra-high Risk for Psychosis and the Association With Clinical Symptoms and Cognition, Biol. Psychiatry Cogn. Neurosci. Neuroimaging. 5 (2020) 569–579. https://doi.org/10.1016/j.bpsc.2019.12.005.

[165] N.R. Bolo, A.M. Jacobson, G. Musen, M.S. Keshavan, D.C. Simonson, Acute Hyperglycemia Increases Brain Pregenual Anterior Cingulate Cortex Glutamate Concentrations in Type 1 Diabetes, Diabetes. 69 (2020) 1528–1539. https://doi.org/10.2337/db19-0936.

[166] A.P. Prescot, S.R. Miller, G. Ingenito, R.S. Huber, D.G. Kondo, P.F. Renshaw, In Vivo Detection of CPP-115 Target Engagement in Human Brain, Neuropsychopharmacology. 43 (2018) 646–654. https://doi.org/10.1038/npp.2017.156.

[167] S.A. Wijtenburg, J. Near, S.A. Korenic, F.E. Gaston, H.J. Chen, M. Mikkelsen, S. Chen, P. Kochunov, L.E. Hong, L.M. Rowland, Comparing the reproducibility of commonly used magnetic resonance spectroscopy techniques to quantify cerebral glutathione, J. Magn. Reson. IMAGING. 49 (2019) 176–183. https://doi.org/10.1002/jmri.26046.

[168] S. Younis, A. Hougaard, C.E. Christensen, M.B. Vestergaard, E.T. Petersen, V.O. Boer, O.B. Paulson, M. Ashina, A. Marsman, H.B.W. Larsson, Feasibility of Glutamate and GABA Detection in Pons and Thalamus at 3T and 7T by Proton Magnetic Resonance Spectroscopy, Front. Neurosci. 14 (2020). https://doi.org/10.3389/fnins.2020.559314.

[169] K.C. Yang, B.H. Yang, J.F. Lirng, M.N. Liu, L.Y. Hu, Y.J. Liou, L.A. Chan, Y.H. Chou, Interaction of dopamine transporter and metabolite ratios underpinning the cognitive dysfunction in patients with carbon monoxide poisoning: A combined SPECT and MRS study, Neurotoxicology. 82 (2021) 26–34. https://doi.org/10.1016/j.neuro.2020.11.002.

[170] L. Simani, S. Raminfard, M. Asadollahi, M. Roozbeh, F. Ryan, M. Rostami, Neurochemicals of limbic system and thalamofrontal cortical network: Are they different between patients with idiopathic generalized epilepsy and psychogenic nonepileptic seizure?, Epilepsy Behav. 112 (2020) 107480. https://doi.org/10.1016/j.yebeh.2020.107480.

[171] V. Tiwari, Z. An, Y. Wang, C. Choi, Distinction of the GABA 2.29 ppm resonance using triple refocusing at 3 T in vivo, Magn. Reson. Med. 80 (2018) 1307–1319. https://doi.org/10.1002/mrm.27142.

[172] M. Mikkelsen, D.L. Rimbault, P.B. Barker, P.K. Bhattacharyya, M.K. Brix, P.F. Buur, K.M. Cecil, K.L. Chan, D.Y.T. Chen, A.R. Craven, K. Cuypers, M. Dacko, N.W. Duncan, U. Dydak, D.A. Edmondson, G. Ende, L. Ersland, M.A. Forbes, F. Gao, I. Greenhouse, A.D. Harris, N. He, S. Heba, N. Hoggard, T.W. Hsu, J.F.A. Jansen, A. Kangarlu, T. Lange, R.M. Lebel, Y. Li, C.Y.E. Lin, J.K. Liou, J.F. Lirng, F. Liu, J.R. Long, R. Ma, C. Maes, M. Moreno-Ortega, S.O. Murray, S. Noah, R. Noeske, M.D. Noseworthy, G. Oeltzschner, E.C. Porges, J.J. Prisciandaro, N.A.J. Puts, T.P.L. Roberts, M. Sack, N. Sailasuta, M.G. Saleh, M.P. Schallmo, N. Simard, D. Stoffers, S.P. Swinnen, M. Tegenthoff, P. Truong, G. Wang, I.D. Wilkinson, H.J. Wittsack, A.J. Woods, H. Xu, F. Yan, C. Zhang, V. Zipunnikov, H.J. Zöllner, R.A.E. Edden, Big GABA II: Water-referenced edited MR spectroscopy at 25 research sites, Neuroimage. 191 (2019) 537–548. https://doi.org/10.1016/j.neuroimage.2019.02.059.

[173] Q.X. Wu, C. Qi, J. Long, Y.H. Liao, X.Y. Wang, A. Xie, J.B. Liu, W. Hao, Y.Y. Tang, B.Z. Yang, T.Q. Liu, J.S. Tang, Metabolites Alterations in the Medial Prefrontal Cortex of Methamphetamine Users in Abstinence: A H-1 MRS Study, Front. PSYCHIATRY. 9 (2018). https://doi.org/10.3389/fpsyt.2018.00478.

[174] E. Plitman, S. Chavez, S. Nakajima, Y. Iwata, J.K. Chung, F. Caravaggio, J. Kim, Y. Alshehri, M.M. Chakravarty, V. De Luca, G. Remington, P. Gerretsen, A. Graff-Guerrero, Striatal neurometabolite levels in patients with schizophrenia undergoing long-term antipsychotic treatment: A proton magnetic resonance spectroscopy and reliability study, Psychiatry Res. - Neuroimaging. 273 (2018) 16–24. https://doi.org/10.1016/j.pscychresns.2018.01.004.

[175] J.D. Xia, F. Chen, Q.J. Zhang, Y.M. Wang, Y.T. Dai, N.H. Song, Z.J. Wang, B. Zhang, J. Yang, Abnormal Thalamic Metabolism in Patients With Lifelong Premature Ejaculation, J. Sex. Med. 18 (2021) 275–283. https://doi.org/10.1016/j.jsxm.2020.11.014.

[176] E.C. Onwordi, T. Whitehurst, A. Mansur, B. Statton, A. Berry, M. Quinlan, D.P. O’Regan, M. Rogdaki, T.R. Marques, E.A. Rabiner, R.N. Gunn, A.C. Vernon, S. Natesan, O.D. Howes, The relationship between synaptic density marker SV2A, glutamate and N-acetyl aspartate levels in healthy volunteers and schizophrenia: a multimodal PET and magnetic resonance spectroscopy brain imaging study, Transl. Psychiatry. 11 (2021) 1–9. https://doi.org/10.1038/s41398-021-01515-3.

[177] D. Hong, S.R. Rankouhi, J.W. Thielen, J.J.A. van Asten, D.G. Norris, A comparison of sLASER and MEGA-sLASER using simultaneous interleaved acquisition for measuring GABA in the human brain at 7T, PLoS One. 14 (2019). https://doi.org/10.1371/journal.pone.0223702.

[178] C. Jiménez-Espinoza, F. Marcano Serrano, J.L. González-Mora, N-acetylaspartyl-glutamate metabolism in the cingulated cortices as a biomarker of the etiology in asd: A1 h-mrs model, Molecules. 26 (2021). https://doi.org/10.3390/molecules26030675.

[179] S. Lai, S. Zhong, X. Liao, Y. Wang, J. Huang, S. Zhang, Y. Sun, H. Zhao, Y. Jia, Biochemical abnormalities in basal ganglia and executive dysfunction in acute- and euthymic-episode patients with bipolar disorder: A proton magnetic resonance spectroscopy study, J. Affect. Disord. 225 (2018) 108–116. https://doi.org/10.1016/j.jad.2017.07.036.

[180] D.E. Harper, E. Ichesco, A. Schrepf, M. Halvorson, T. Puiu, D.J. Clauw, R.E. Harris, S.E. Harte, M.R. Network, Relationships between brain metabolite levels, functional connectivity, and negative mood in urologic chronic pelvic pain syndrome patients compared to controls: A MAPP research network study, NEUROIMAGE-CLINICAL. 17 (2018) 570–578. https://doi.org/10.1016/j.nicl.2017.11.014.

[181] Y.H. Jung, H. Kim, D. Lee, J.Y. Lee, J.Y. Moon, S.H. Choi, D.H. Kang, Dysfunctional energy metabolisms in fibromyalgia compared with healthy subjects, Mol. Pain. 17 (2021). https://doi.org/10.1177/17448069211012833.

[182] T. Gradinger, M. Sack, V. Cardinale, M. Thiacourt, U. Baumgartner, C. Schmahl, G. Ende, The glutamate to g-Aminobutyric acid ratio in the posterior insula is associated with pain perception in healthy women but not in women with borderline personality disorder, Pain. 160 (2019) 2487–2496. https://doi.org/10.1097/j.pain.0000000000001641.

[183] J.N. Li, X.L. Liu, L. Li, Prefrontal GABA and glutamate levels correlate with impulsivity and cognitive function of prescription opioid addicts: A H-1-magnetic resonance spectroscopy study, PSYCHIATRY Clin. Neurosci. 74 (2020) 77–83. https://doi.org/10.1111/pcn.12940.

[184] T. Borbath, S. Murali-Manohar, A.M. Wright, A. Henning, In vivo characterization of downfield peaks at 9.4 T: T(2)relaxation times, quantification, pH estimation, and assignments, Magn. Reson. Med. (n.d.). https://doi.org/10.1002/mrm.28442.

[185] F. Jabbari-Zadeh, B. Cao, J.A. Stanley, Y. Liu, M.J. Wu, J. Tannous, M. Lopez, M. Sanches, B. Mwangi, G.B. Zunta-Soares, J.C. Soares, Evidence of altered metabolism of cellular membranes in bipolar disorder comorbid with post-traumatic stress disorder, J. Affect. Disord. 289 (2021) 81–87. https://doi.org/10.1016/j.jad.2021.04.011.

[186] K.L. Benson, R. Bottary, L. Schoerning, L. Baer, A. Gonenc, J.E. Jensen, J.W. Winkelman, 1H MRS Measurement of Cortical GABA and Glutamate in Primary Insomnia and Major Depressive Disorder: Relationship to Sleep Quality and Depression Severity, J. Affect. Disord. 274 (2020) 624–631. https://doi.org/10.1016/j.jad.2020.05.026.

[187] Y. Akiyama, R. Yokoyama, H. Takashima, Y. Kawata, M. Arihara, R. Chiba, Y. Kimura, T. Mikami, N. Mikuni, Accumulation of macromolecules in idiopathic normal pressure hydrocephalus, Neurol. Med. Chir. (Tokyo). 61 (2021) 211–218. https://doi.org/10.2176/nmc.oa.2020-0274.

[188] A.M. Wang, S. Pradhan, J.M. Coughlin, A. Trivedi, S.L. DuBois, J.L. Crawford, T.W. Sedlak, F.C. Nucifora, G. Nestadt, L.G. Nucifora, D.J. Schretlen, A. Sawa, P.B. Barker, Assessing Brain Metabolism With 7-T Proton Magnetic Resonance Spectroscopy in Patients With First-Episode Psychosis, JAMA PSYCHIATRY. 76 (2019) 314–323. https://doi.org/10.1001/jamapsychiatry.2018.3637.

[189] J.R. Bustillo, T. Jones, C. Qualls, L. Chavez, D. Lin, R.K. Lenroot, C. Gasparovic, Proton magnetic resonance spectroscopic imaging of gray and white matter in bipolar-I and schizophrenia, J. Affect. Disord. 246 (2019) 745–753. https://doi.org/10.1016/j.jad.2018.12.064.

[190] T. Bell, E.S. Boudes, R.S. Loo, G.J. Barker, D.J. Lythgoe, R.A.E. Edden, R.M. Lebel, M. Wilson, A.D. Harris, In vivo Glx and Glu measurements from GABA-edited MRS at 3 T, NMR Biomed. 34 (2021). https://doi.org/10.1002/nbm.4245.

[191] S.A. Korenic, E.A. Klingaman, E.M. Wickwire, F.E. Gaston, H.J. Chen, S.A. Wijtenburg, L.M. Rowland, Sleep quality is related to brain glutamate and symptom severity in schizophrenia, J. Psychiatr. Res. 120 (2020) 14–20. https://doi.org/10.1016/j.jpsychires.2019.10.006.

[192] N.L. Mason, E.L. Theunissen, N. Hutten, D.H.Y. Tse, S.W. Toennes, P. Stiers, J.G. Ramaekers, Cannabis induced increase in striatal glutamate associated with loss of functional corticostriatal connectivity, Eur. Neuropsychopharmacol. 29 (2019) 247–256. https://doi.org/10.1016/j.euroneuro.2018.12.003.

[193] M. Shimizu, Y. Suzuki, K. Yamada, S. Ueki, M. Watanabe, H. Igarashi, T. Nakada, Maturational decrease of glutamate in the human cerebral cortex from childhood to young adulthood: a H-1-MR spectroscopy study, Pediatr. Res. 82 (2017) 749–752. https://doi.org/10.1038/pr.2017.101.

[194] A. Lind, C.J. Boraxbekk, E.T. Petersen, O.B. Paulson, H.R. Siebner, A. Marsman, Regional Myo-Inositol, Creatine, and Choline Levels Are Higher at Older Age and Scale Negatively with Visuospatial Working Memory: A Cross-Sectional Proton MR Spectroscopy Study at 7 Tesla on Normal Cognitive Ageing, J. Neurosci. 40 (2020) 8149–8159. https://doi.org/10.1523/JNEUROSCI.2883-19.2020.

[195] M.C. Ferland, J.M. Therrien-Blanchet, G. Lefebvre, G. Klees-Themens, S. Proulx, H. Théoret, Longitudinal assessment of 1H-MRS (GABA and Glx) and TMS measures of cortical inhibition and facilitation in the sensorimotor cortex, Exp. Brain Res. 237 (2019) 3461–3474. https://doi.org/10.1007/s00221-019-05691-z.

[196] E. Lyros, A. Ragoschke-Schumm, P. Kostopoulos, A. Sehr, M. Backens, S. Kalampokini, Y. Decker, M. Lesmeister, Y. Liu, W. Reith, K. Fassbender, Normal brain aging and Alzheimer’s disease are associated with lower cerebral pH: an in vivo histidine 1H-MR spectroscopy study, Neurobiol. Aging. 87 (2020) 60–69. https://doi.org/10.1016/j.neurobiolaging.2019.11.012.

[197] B.R. Godlewska, A. Minichino, U. Emir, I. Angelescu, B. Lennox, M. Micunovic, O. Howes, P.J. Cowen, Brain glutamate concentration in men with early psychosis: a magnetic resonance spectroscopy case-control study at 7 T, Transl. Psychiatry. 11 (2021) 367. https://doi.org/10.1038/s41398-021-01477-6.

[198] A. Egerton, B. V. Broberg, N. Van Haren, K. Merritt, G.J. Barker, D.J. Lythgoe, R. Perez-Iglesias, L. Baandrup, S.W. Düring, K. V. Sendt, J.M. Stone, E. Rostrup, I.E. Sommer, B. Glenthøj, R.S. Kahn, P. Dazzan, P. McGuire, Response to initial antipsychotic treatment in first episode psychosis is related to anterior cingulate glutamate levels: a multicentre 1 H-MRS study (OPTiMiSE), Mol. Psychiatry. 23 (2018) 2145–2155. https://doi.org/10.1038/s41380-018-0082-9.

[199] Y.H. Jung, H. Kim, D. Lee, J.Y. Lee, W.J. Lee, J.Y. Moon, S.H. Choi, D.H. Kang, Abnormal neurometabolites in fibromyalgia patients: Magnetic resonance spectroscopy study, Mol. Pain. 17 (2021). https://doi.org/10.1177/1744806921990946.

[200] R. Wang, Q. Fan, Z. Zhang, Y. Chen, Y. Zhu, Y. Li, Anterior thalamic radiation structural and metabolic changes in obsessive-compulsive disorder: A combined DTI-MRS study, Psychiatry Res. - Neuroimaging. 277 (2018) 39–44. https://doi.org/10.1016/j.pscychresns.2018.05.004.

[201] S. Marenco, C. Meyer, J.W. van der Veen, Y. Zhang, R. Kelly, J. Shen, D.R. Weinberger, D. Dickinson, K.F. Berman, Role of gamma-amino-butyric acid in the dorsal anterior cingulate in age-associated changes in cognition, NEUROPSYCHOPHARMACOLOGY. 43 (2018) 2285–2291. https://doi.org/10.1038/s41386-018-0134-5.

[202] F.R. Borgan, S. Jauhar, R.A. McCutcheon, F.S. Pepper, M. Rogdaki, D.J. Lythgoe, O.D. Howes, Glutamate levels in the anterior cingulate cortex in un-medicated first episode psychosis: a proton magnetic resonance spectroscopy study, Sci. Rep. 9 (2019) 1–10. https://doi.org/10.1038/s41598-019-45018-0.

[203] J.B.D. Andrade, F.M. Ferreira, C. Suo, M. Yucel, I. Frydman, M. Monteiro, P. Vigne, L.F. Fontenelle, F. Tovar-Moll, An MRI Study of the Metabolic and Structural Abnormalities in Obsessive-Compulsive Disorder, Front. Hum. Neurosci. 13 (2019). https://doi.org/10.3389/fnhum.2019.00186.

[204] J. Persson, A. Wall, J. Weis, M. Gingnell, G. Antoni, M. Lubberink, R. Bodén, Inhibitory and excitatory neurotransmitter systems in depressed and healthy: A positron emission tomography and magnetic resonance spectroscopy study, Psychiatry Res. - Neuroimaging. 315 (2021). https://doi.org/10.1016/j.pscychresns.2021.111327.

[205] S. Chenji, E. Cox, N. Jaworska, R.M. Swansburg, F.P. MacMaster, Body mass index and variability in hippocampal volume in youth with major depressive disorder, J. Affect. Disord. 282 (2021) 415–425. https://doi.org/10.1016/j.jad.2020.12.176.

[206] O. Al-iedani, J. Arm, K. Ribbons, R. Lea, J. Lechner-Scott, S. Ramadan, Diurnal stability and long-term repeatability of neurometabolites using single voxel 1H magnetic resonance spectroscopy, Eur. J. Radiol. 108 (2018) 107–113. https://doi.org/10.1016/j.ejrad.2018.09.020.

[207] T. Gu, L. Lin, Y. Jiang, J. Chen, R.C.N. D’Arcy, M. Chen, X.W. Song, Acupuncture therapy in treating migraine: results of a magnetic resonance spectroscopy imaging study, J. Pain Res. 11 (2018) 889–900. https://doi.org/10.2147/JPR.S162696.

[208] A. Burger, M.J. Kotze, D.J. Stein, S.J. van Rensburg, F.M. Howells, The relationship between measurement of in vivo brain glutamate and markers of iron metabolism: A proton magnetic resonance spectroscopy study in healthy adults, Eur. J. Neurosci. 51 (2020) 984–990. https://doi.org/10.1111/ejn.14583.

[209] D.F. Hermens, S.N. Hatton, R.S.C. Lee, S.L. Naismith, S.L. Duffy, G. Paul Amminger, M. Kaur, E.M. Scott, J. Lagopoulos, I.B. Hickie, In vivo imaging of oxidative stress and fronto-limbic white matter integrity in young adults with mood disorders, Eur. Arch. Psychiatry Clin. Neurosci. 268 (2018) 145–156. https://doi.org/10.1007/s00406-017-0788-8.

[210] M.G. Soeiro-de-Souza, M.C.G. Otaduy, R. Machado-Vieira, R.A. Moreno, F.G. Nery, C. Leite, B. Lafer, Lithium-associated anterior cingulate neurometabolic profile in euthymic Bipolar I disorder: A 1H-MRS study, J. Affect. Disord. 241 (2018) 192–199. https://doi.org/10.1016/j.jad.2018.08.039.

[211] L.L. Gramegna, S. Evangelisti, L. Di Vito, C. La Morgia, A. Maresca, L. Caporali, G. Amore, L. Talozzi, C. Bianchini, C. Testa, D.N. Manners, I. Cortesi, M.L. Valentino, R. Liguori, V. Carelli, C. Tonon, R. Lodi, Brain MRS correlates with mitochondrial dysfunction biomarkers in MELAS-associated mtDNA mutations, Ann. Clin. Transl. Neurol. 8 (2021) 1200–1211. https://doi.org/10.1002/acn3.51329.

[212] S. Younis, A. Hougaard, C.E. Christensen, M.B. Vestergaard, E.T. Petersen, O.B. Paulson, H.B.W. Larsson, M. Ashina, Effects of sildenafil and calcitonin gene-related peptide on brainstem glutamate levels: a pharmacological proton magnetic resonance spectroscopy study at 3.0 T, J. Headache Pain. 19 (2018). https://doi.org/10.1186/s10194-018-0870-2.

[213] S. Zhong, Y. Wang, S. Lai, T. Liu, X. Liao, G. Chen, Y. Jia, Associations between executive function impairment and biochemical abnormalities in bipolar disorder with suicidal ideation, J. Affect. Disord. 241 (2018) 282–290. https://doi.org/10.1016/j.jad.2018.08.031.

[214] S.A. Rafique, J.K.E. Steeves, Assessing differential effects of single and accelerated low-frequency rTMS to the visual cortex on GABA and glutamate concentrations, Brain Behav. 10 (2020) 1–18. https://doi.org/10.1002/brb3.1845.

[215] K. Ryan, K. Wawrzyn, J.S. Gati, B.A. Chronik, D. Wong, N. Duggal, R. Bartha, 1H MR spectroscopy of the motor cortex immediately following transcranial direct current stimulation at 7 Tesla, PLoS One. 13 (2018) 1–14. https://doi.org/10.1371/journal.pone.0198053.

[216] F. Borgan, M. Veronese, T. Reis Marques, D.J. Lythgoe, O. Howes, Association between cannabinoid 1 receptor availability and glutamate levels in healthy controls and drug-free patients with first episode psychosis: a multi-modal PET and 1H-MRS study, Eur. Arch. Psychiatry Clin. Neurosci. 271 (2021) 677–687. https://doi.org/10.1007/s00406-020-01191-2.

[217] C. Graf, E.L. MacMillan, E. Fu, T. Harris, A. Traboulsee, I.M. Vavasour, A.L. MacKay, B. Madler, D.K.B. Li, C. Laule, Intra- and inter-site reproducibility of human brain single-voxel proton MRS at 3 T, NMR Biomed. 32 (2019). https://doi.org/10.1002/nbm.4083.

[218] S.Y. Kim, M.J. Kaufman, B.M. Cohen, J.E. Jensen, J.T. Coyle, F. Du, D. Ongur, In Vivo Brain Glycine and Glutamate Concentrations in Patients With First-Episode Psychosis Measured by Echo Time-Averaged Proton Magnetic Resonance Spectroscopy at 4T, Biol. Psychiatry. 83 (2018) 484–491. https://doi.org/10.1016/j.biopsych.2017.08.022.

[219] K.F. Shattuck, J.W. VanMeter, Task-based changes in proton MR spectroscopy signal during configural working memory in human medial temporal lobe, J. Magn. Reson. IMAGING. 47 (2018) 682–691. https://doi.org/10.1002/jmri.25816.

[220] J.R. Bustillo, E.G. Mayer, J. Upston, T. Jones, C. Garcia, S. Sheriff, A. Maudsley, M. Tohen, C. Gasparovic, R. Lenroot, Increased Glutamate Plus Glutamine in the Right Middle Cingulate in Early Schizophrenia but Not in Bipolar Psychosis: A Whole Brain H-1-MRS Study, Front. PSYCHIATRY. 12 (2021). https://doi.org/10.3389/fpsyt.2021.660850.

[221] S. Smesny, J. Große, A. Gussew, K. Langbein, N. Schönfeld, G. Wagner, M. Valente, J.R. Reichenbach, Prefrontal glutamatergic emotion regulation is disturbed in cluster B and C personality disorders – A combined 1H/31P-MR spectroscopic study, J. Affect. Disord. 227 (2018) 688–697. https://doi.org/10.1016/j.jad.2017.10.044.

[222] O.T. Ousdal, A.M. Milde, A.R. Craven, L. Ersland, T. Endestad, A. Melinder, Q.J. Huys, K. Hugdahl, Prefrontal glutamate levels predict altered amygdala-prefrontal connectivity in traumatized youths, Psychol. Med. 49 (2019) 1822–1830. https://doi.org/10.1017/S0033291718002519.

[223] J. Boban, D. Kozic, V. Turkulov, J. Ostojic, R. Semnic, D. Lendak, S. Brkic, HIV-associated neurodegeneration and neuroimmunity: multivoxel MR spectroscopy study in drug-naïve and treated patients, Eur. Radiol. 27 (2017) 4218–4236. https://doi.org/10.1007/s00330-017-4772-5.

[224] F.A. Provenzano, J. Guo, M.M. Wall, X.Y. Feng, H.C. Sigmon, G. Brucato, M.B. First, D.L. Rothman, R.R. Girgis, J.A. Lieberman, S.A. Small, Hippocampal Pathology in Clinical High-Risk Patients and the Onset of Schizophrenia, Biol. Psychiatry. 87 (2020) 234–242. https://doi.org/10.1016/j.biopsych.2019.09.022.

[225] M. Simmonite, J. Carp, B.R. Foerster, L. Ossher, M. Petrou, D.H. Weissman, T.A. Polk, Age-Related Declines in Occipital GABA are Associated with Reduced Fluid Processing Ability, Acad. Radiol. 26 (2019) 1053–1061. https://doi.org/10.1016/j.acra.2018.07.024.

[226] T. Okada, H. Kuribayashi, L.G. Kaiser, Y. Urushibata, N. Salibi, R.T. Seethamraju, S. Ahn, D.H.D. Thuy, K. Fujimoto, T. Isa, Repeatability of proton magnetic resonance spectroscopy of the brain at 7 T: effect of scan time on semi-localized by adiabatic selective refocusing and short-echo time stimulated echo acquisition mode scans and their comparison, Quant. Imaging Med. Surg. 11 (2021) 9–20. https://doi.org/10.21037/qims-20-517.

[227] P. Faulkner, S. Lucini Paioni, P. Kozhuharova, N. Orlov, D.J. Lythgoe, Y. Daniju, E. Morgenroth, H. Barker, P. Allen, Daily and intermittent smoking are associated with low prefrontal volume and low concentrations of prefrontal glutamate, creatine, myo-inositol, and N-acetylaspartate, Addict. Biol. 26 (2021) 1–11. https://doi.org/10.1111/adb.12986.

[228] T. Bell, M. Lindner, A. Langdon, P.G. Mullins, A. Christakou, Regional Striatal Cholinergic Involvement in Human Behavioral Flexibility, J. Neurosci. 39 (2019) 5740–5749. https://doi.org/10.1523/JNEUROSCI.2110-18.2019.

[229] P.W. Chiu, S.S.Y. Lui, K.S.Y. Hung, R.C.K. Chan, Q. Chan, P.C. Sham, E.F.C. Cheung, H.K.F. Mak, In vivo gamma-aminobutyric acid and glutamate levels in people with first-episode schizophrenia: A proton magnetic resonance spectroscopy study, Schizophr. Res. 193 (2018) 295–303. https://doi.org/10.1016/j.schres.2017.07.021.

[230] B. Cao, J.A. Stanley, I.C. Passos, B. Mwangi, S. Selvaraj, G.B. Zunta-Soares, J.C. Soares, Elevated Choline-Containing Compound Levels in Rapid Cycling Bipolar Disorder, NEUROPSYCHOPHARMACOLOGY. 42 (2017) 2252–2258. https://doi.org/10.1038/npp.2017.39.

[231] I. Betina Ip, U.E. Emir, A.J. Parker, J. Campbell, H. Bridge, Comparison of neurochemical and BOLD signal contrast response functions in the human visual cortex, J. Neurosci. 39 (2019) 7968–7975. https://doi.org/10.1523/JNEUROSCI.3021-18.2019.

[232] J.W. Evans, N. Lally, L. An, N.Z. Li, A.C. Nugent, D. Banerjee, S.L. Snider, J. Shen, J.P. Roiser, C.A. Zarate, 7T H-1-MRS in major depressive disorder: a Ketamine Treatment Study, NEUROPSYCHOPHARMACOLOGY. 43 (2018) 1908–1914. https://doi.org/10.1038/s41386-018-0057-1.

[233] Y. Boillat, L. Xin, W. van der Zwaag, R. Gruetter, Metabolite concentration changes associated with positive and negative BOLD responses in the human visual cortex: A functional MRS study at 7 Tesla, J. Cereb. Blood Flow Metab. 40 (2020) 488–500. https://doi.org/10.1177/0271678X19831022.

[234] M.G. Saleh, M. Mikkelsen, G. Oeltzschner, K.L. Chan, A. Berrington, P.B. Barker, R.A.E. Edden, Simultaneous editing of GABA and glutathione at 7T using semi-LASER localization, Magn. Reson. Med. 80 (2018) 474–479. https://doi.org/10.1002/mrm.27044.

[235] M.A. Monnig, A.J. Woods, E. Walsh, C.M. Martone, J. Blumenthal, P.M. Monti, R.A. Cohen, Cerebral Metabolites on the Descending Limb of Acute Alcohol: A Preliminary 1H MRS Study, Alcohol Alcohol. 54 (2019) 487–496. https://doi.org/10.1093/alcalc/agz062.

[236] M. Karczewska-Kupczewska, A. Nikolajuk, R. Filarski, R. Majewski, E. Tarasow, Intralipid/Heparin Infusion Alters Brain Metabolites Assessed With H-1-MRS Spectroscopy in Young Healthy Men, J. Clin. Endocrinol. Metab. 103 (2018) 2563–2570. https://doi.org/10.1210/jc.2018-00107.

[237] T. Kameda, S. Fukui, R. Tominaga, M. Sekiguchi, N. Iwashita, K. Ito, S. Tanaka-Mizuno, S.I. Konno, Brain metabolite changes in the anterior cingulate cortex of chronic low back pain patients and correlations between metabolites and psychological state, Clin. J. Pain. 34 (2018) 657–663. https://doi.org/10.1097/AJP.0000000000000583.

[238] P. Bednařík, I. Tkáč, F. Giove, L.E. Eberly, D.K. Deelchand, F.R. Barreto, S. Mangia, Neurochemical responses to chromatic and achromatic stimuli in the human visual cortex, J. Cereb. Blood Flow Metab. 38 (2018) 347–359. https://doi.org/10.1177/0271678X17695291.

[239] M. Terpstra, C. Torkelson, U. Emir, J.S. Hodges, S. Raatz, Noninvasive quantification of human brain antioxidant concentrations after an intravenous bolus of vitamin C, NMR Biomed. 24 (2011) 521–528. https://doi.org/10.1002/nbm.1619.

[240] J.E. Siegel-Ramsay, L. Romaniuk, H.C. Whalley, N. Roberts, H. Branigan, A.C. Stanfield, S.M. Lawrie, M.R. Dauvermann, Glutamate and functional connectivity - support for the excitatory-inhibitory imbalance hypothesis in autism spectrum disorders, Psychiatry Res. - Neuroimaging. 313 (2021) 111302. https://doi.org/10.1016/j.pscychresns.2021.111302.

[241] I. Savic, MRS Shows Regionally Increased Glutamate Levels among Patients with Exhaustion Syndrome Due to Occupational Stress, Cereb. Cortex. 30 (2020) 3759–3770. https://doi.org/10.1093/cercor/bhz340.

[242] S. Smesny, D. Berberich, A. Gussew, N. Schönfeld, K. Langbein, M. Walther, J.R. Reichenbach, Alterations of neurometabolism in the dorsolateral prefrontal cortex and thalamus in transition to psychosis patients change under treatment as usual – A two years follow-up 1H/31P-MR-spectroscopy study, Schizophr. Res. 228 (2021) 7–18. https://doi.org/10.1016/j.schres.2020.11.063.

[243] A.L. Peek, A.M. Leaver, S. Foster, G. Oeltzschner, N.A. Puts, G. Galloway, M. Sterling, K. Ng, K. Refshauge, M.E.R. Aguila, T. Rebbeck, Increased GABA+ in People With Migraine, Headache, and Pain Conditions- A Potential Marker of Pain, J. Pain. 22 (2021) 1631–1645. https://doi.org/10.1016/j.jpain.2021.06.005.

[244] I.I. Kirov, M. Sollberger, M.S. Davitz, L. Glodzik, B.J. Soher, J.S. Babb, A.U. Monsch, A. Gass, O. Gonen, Global brain volume and N-acetyl-aspartate decline over seven decades of normal aging, Neurobiol. Aging. 98 (2021) 42–51. https://doi.org/10.1016/j.neurobiolaging.2020.10.024.

[245] Y. Zhang, E. Taub, C. Mueller, J. Younger, G. Uswatte, T.P. DeRamus, D.C. Knight, Reproducibility of whole-brain temperature mapping and metabolite quantification using proton magnetic resonance spectroscopy, NMR Biomed. 33 (2020) 1–13. https://doi.org/10.1002/nbm.4313.

[246] L.A. Jelen, S. King, C.M. Horne, D.J. Lythgoe, A.H. Young, J.M. Stone, Functional magnetic resonance spectroscopy in patients with schizophrenia and bipolar affective disorder: Glutamate dynamics in the anterior cingulate cortex during a working memory task, Eur. Neuropsychopharmacol. 29 (2019) 222–234. https://doi.org/10.1016/j.euroneuro.2018.12.005.

[247] J.P. Hegarty, D.J. Weber, C.M. Cirstea, D.Q. Beversdorf, Cerebro-Cerebellar Functional Connectivity is Associated with Cerebellar Excitation-Inhibition Balance in Autism Spectrum Disorder, J. Autism Dev. Disord. 48 (2018) 3460–3473. https://doi.org/10.1007/s10803-018-3613-y.

[248] J.J. Prisciandaro, J.P. Schacht, A.P. Prescot, P.F. Renshaw, T.R. Brown, R.F. Anton, Brain Glutamate, GABA, and Glutamine Levels and Associations with Recent Drinking in Treatment-Naive Individuals with Alcohol Use Disorder Versus Light Drinkers, Alcohol. Exp. Res. 43 (2019) 221–226. https://doi.org/10.1111/acer.13931.

[249] A. Garkowski, B. Kubas, M. Hładuński, J. Zajkowska, O. Zajkowska, D. Jurgilewicz, R. Zawadzki, E. Garkowska, S. Pancewicz, U. Łebkowska, Neuronal loss or dysfunction in patients with early Lyme neuroborreliosis: a proton magnetic resonance spectroscopy study of the brain, J. Neurol. 266 (2019) 1937–1943. https://doi.org/10.1007/s00415-019-09359-0.

[250] M.G. Soeiro-de-Souza, M.C.G. Otaduy, R. Machado-Vieira, R.A. Moreno, F.G. Nery, C. Leite, B. Lafer, Anterior Cingulate Cortex Glutamatergic Metabolites and Mood Stabilizers in Euthymic Bipolar I Disorder Patients: A Proton Magnetic Resonance Spectroscopy Study, Biol. Psychiatry Cogn. Neurosci. Neuroimaging. 3 (2018) 985–991. https://doi.org/10.1016/j.bpsc.2018.02.007.

[251] Q.Y. Tan, H.Q. Sun, W.N. Wang, X.T. Wu, N.Y. Hao, X.R. Su, X.B. Yang, S.M. Zhang, J.K. Su, Q. Yue, Q.Y. Gong, Quantitative MR spectroscopy reveals metabolic changes in the dorsolateral prefrontal cortex of patients with temporal lobe epilepsy, Eur. Radiol. 28 (2018) 4496–4503. https://doi.org/10.1007/s00330-018-5443-x.

[252] X. Liu, S. Zhong, Z. Li, J. Chen, Y. Wang, S. Lai, H. Miao, Y. Jia, Serum copper and zinc levels correlate with biochemical metabolite ratios in the prefrontal cortex and lentiform nucleus of patients with major depressive disorder, Prog. Neuro-Psychopharmacology Biol. Psychiatry. 99 (2020) 109828. https://doi.org/10.1016/j.pnpbp.2019.109828.

[253] Y. Iwata, S. Nakajima, E. Plitman, F. Caravaggio, J. Kim, P. Shah, W. Mar, S. Chavez, V. De Luca, M. Mimura, G. Remington, P. Gerretsen, A. Graff-Guerrero, Glutamatergic Neurometabolite Levels in Patients With Ultra-Treatment-Resistant Schizophrenia: A Cross-Sectional 3T Proton Magnetic Resonance Spectroscopy Study, Biol. Psychiatry. 85 (2019) 596– 605. https://doi.org/10.1016/j.biopsych.2018.09.009.

[254] E.C. Wiegers, H.M. Rooijackers, J.J.A. van Asten, C.J. Tack, A. Heerschap, B.E. de Galan, M. van der Graaf, Elevated brain glutamate levels in type 1 diabetes: correlations with glycaemic control and age of disease onset but not with hypoglycaemia awareness status, Diabetologia. 62 (2019) 1065–1073. https://doi.org/10.1007/s00125-019-4862-9.

[255] B.P. Brennan, R. Admon, C. Perriello, E.M. LaFlamme, A.J. Athey, D.A. Pizzagalli, J.I. Hudson, H.G. Pope, J.E. Jensen, Acute change in anterior cingulate cortex GABA, but not glutamine/glutamate, mediates antidepressant response to citalopram, PSYCHIATRY Res. 269 (2017) 9–16. https://doi.org/10.1016/j.pscychresns.2017.08.009.

[256] D.K. Deelchand, M. Marjańska, J.S. Hodges, M. Terpstra, Sensitivity and specificity of human brain glutathione concentrations measured using short-TE 1H MRS at 7 T, NMR Biomed. 29 (2016) 600–606. https://doi.org/10.1002/nbm.3507.

[257] P. Hnilicová, E. Kantorová, H. Poláček, M. Grendár, M. Bittšanský, D. Čierny, Š. Sivák, K. Zeleňák, J. Lehotský, D. Dobrota, E. Kurča, Altered hypothalamic metabolism in early multiple sclerosis – MR spectroscopy study, J. Neurol. Sci. 407 (2019). https://doi.org/10.1016/j.jns.2019.116458.

[258] W. Wang, X. Wu, X. Su, H. Sun, Q. Tan, S. Zhang, L. Lu, H. Gao, W. Liu, X. Yang, D. Zhou, G.J. Kemp, Q. Yue, Q. Gong, Metabolic alterations of the dorsolateral prefrontal cortex in sleep-related hypermotor epilepsy: A proton magnetic resonance spectroscopy study, J. Neurosci. Res. 99 (2021) 2657–2668. https://doi.org/10.1002/jnr.24866.

[259] G. Barbagallo, G. Arabia, M. Morelli, R. Nisticò, F. Novellino, M. Salsone, F. Rocca, A. Quattrone, M. Caracciolo, U. Sabatini, A. Cherubini, A. Quattrone, Thalamic neurometabolic alterations in tremulous Parkinson’s disease: A preliminary proton MR spectroscopy study, Park. Relat. Disord. 43 (2017) 78–84. https://doi.org/10.1016/j.parkreldis.2017.07.028.

[260] E.E.A. Elmaki, T. Gong, D.M. Nkonika, G. Wang, Examining alterations in GABA concentrations in the basal ganglia of patients with Parkinson’s disease using MEGA-PRESS MRS, Jpn. J. Radiol. 36 (2018) 194–199. https://doi.org/10.1007/s11604-017-0714-z.

[261] D. Shukla, P.K. Mandal, M. Tripathi, G. Vishwakarma, R. Mishra, K. Sandal, Quantitation of in vivo brain glutathione conformers in cingulate cortex among age-matched control, MCI, and AD patients using MEGA-PRESS, Hum. Brain Mapp. 41 (2020) 194–217. https://doi.org/10.1002/hbm.24799.

[262] B. Schmitz, H. Pflugrad, A.B. Tryc, H. Lanfermann, E. Jäckel, H. Schrem, J. Beneke, H. Barg-Hock, J. Klempnauer, K. Weissenborn, X.Q. Ding, Brain metabolic alterations in patients with long-term calcineurin inhibitor therapy after liver transplantation, Aliment. Pharmacol. Ther. 49 (2019) 1431–1441. https://doi.org/10.1111/apt.15256.

[263] M. Marjańska, J. Riley McCarten, J.S. Hodges, L.S. Hemmy, M. Terpstra, Distinctive Neurochemistry in Alzheimer’s Disease via 7 T in Vivo Magnetic Resonance Spectroscopy, J. Alzheimer’s Dis. 68 (2019) 559–569. https://doi.org/10.3233/JAD-180861.

[264] E.J. Mellen, D.G. Harper, C. Ravichandran, E. Jensen, M. Silveri, B.P. Forester, Lamotrigine Therapy and Biomarkers of Cerebral Energy Metabolism in Older Age Bipolar Depression, Am. J. Geriatr. Psychiatry. 27 (2019) 783–793. https://doi.org/10.1016/j.jagp.2019.02.017.

[265] L. Lu, J. Wang, L. Zhang, Z. Zhang, L. Ni, R. Qi, X. Kong, M. Lu, M.U. Sami, K. Xu, G. Lu, Disrupted metabolic and functional connectivity patterns of the posterior cingulate cortex in cirrhotic patients: A study combining magnetic resonance spectroscopy and resting-state functional magnetic resonance imaging, Neuroreport. 29 (2018) 993–1000. https://doi.org/10.1097/WNR.0000000000001063.

[266] S.C. Craciunas, M.R. Gorgan, B. Ianosi, P. Lee, J. Burris, C.M. Cirstea, Remote motor system metabolic profile and surgery outcome in cervical spondylotic myelopathy, J. Neurosurg. Spine. 26 (2017) 668–678. https://doi.org/10.3171/2016.10.SPINE16479.

[267] G.S. Smith, G. Oeltzschner, N.F. Gould, J.M.S. Leoutsakos, N. Nassery, J.H. Joo, M.A. Kraut, R.A.E. Edden, P.B. Barker, S.A. Wijtenburg, L.M. Rowland, C.I. Workman, Neurotransmitters and Neurometabolites in Late-Life Depression: A Preliminary Magnetic Resonance Spectroscopy Study at 7T, J. Affect. Disord. 279 (2021) 417–425. https://doi.org/10.1016/j.jad.2020.10.011.

[268] L. Mazuel, C. Chassain, B. Jean, B. Pereira, A. Cladière, C. Speziale, F. Durif, Proton MR spectroscopy for diagnosis and evaluation of treatment efficacy in Parkinson disease, Radiology. 278 (2016) 505–513. https://doi.org/10.1148/radiol.2015142764.

[269] M. Mitolo, M. Stanzani-Maserati, D.N. Manners, S. Capellari, C. Testa, L. Talozzi, R. Poda, F. Oppi, S. Evangelisti, L.L. Gramegna, S. Magarelli, R. Pantieri, R. Liguori, R. Lodi, C. Tonon, The combination of metabolic posterior cingulate cortical abnormalities and structural asymmetries improves the differential diagnosis between primary progressive aphasia and alzheimer’s disease, J. Alzheimer’s Dis. 82 (2021) 1467–1473. https://doi.org/10.3233/JAD-210211.

[270] Y.L. Song, T. Gong, Y.Y. Xiang, M. Mikkelsen, G.B. Wang, R.A.E. Edden, Single-dose L-dopa increases upper brainstem GABA in Parkinsons disease: A preliminary study, J. Neurol. Sci. 422 (2021). https://doi.org/10.1016/j.jns.2021.117309.

[271] A.D. Seger, E. Farrher, C.E.J. Doppler, A. Gogishvili, W.A. Worthoff, C.P. Filss, M.T. Barbe, F. Holtbernd, N.J. Shah, G.R. Fink, M. Sommerauer, Putaminal y-Aminobutyric Acid Modulates Motor Response to Dopaminergic Therapy in Parkinson’s Disease, Mov. Disord. 36 (2021) 2187–2192. https://doi.org/10.1002/mds.28674.

[272] G. Oeltzschner, S.A. Wijtenburg, M. Mikkelsen, R.A.E. Edden, P.B. Barker, J.H. Joo, J.M.S. Leoutsakos, L.M. Rowland, C.I. Workman, G.S. Smith, Neurometabolites and associations with cognitive deficits in mild cognitive impairment: a magnetic resonance spectroscopy study at 7 Tesla, Neurobiol. Aging. 73 (2019) 211–218. https://doi.org/10.1016/j.neurobiolaging.2018.09.027.

[273] B. Pesch, S. Casjens, D. Woitalla, S. Dharmadhikari, D.A. Edmondson, M.A.S. Zella, M. Lehnert, A. Lotz, L. Herrmann, S. Muhlack, P. Kraus, C.L. Yeh, B. Glaubitz, T. Schmidt-Wilcke, R. Gold, C. van Thriel, T. Bruning, L. Tonges, U. Dydak, Impairment of Motor Function Correlates with Neurometabolite and Brain Iron Alterations in Parkinson’s Disease, CELLS. 8 (2019). https://doi.org/10.3390/cells8020096.

[274] L.H. Chen, J.Y. Shi, T.X. Zou, L. Zhang, Y. Gou, Y. Lin, H.J. Chen, Disturbance of thalamic metabolism and its association with regional neural dysfunction and cognitive impairment in minimal hepatic encephalopathy, Eur. J. Radiol. 131 (2020) 109252. https://doi.org/10.1016/j.ejrad.2020.109252.

[275] T.T. Tran, K. Wei, S. Cole, E. Mena, M. Csete, K.S. King, Brain MR Spectroscopy Markers of Encephalopathy Due to Nonalcoholic Steatohepatitis, J. Neuroimaging. 30 (2020) 697–703. https://doi.org/10.1111/jon.12728.

[276] S. Chaudhary, S.S. Kumaran, V. Goyal, M. Kalaivani, G.S. Kaloiya, R. Sagar, N. Mehta, A.K. Srivastava, N.R. Jagannathan, Frontal lobe metabolic alterations characterizing Parkinson’s disease cognitive impairment, Neurol. Sci. 42 (2021) 1053–1064. https://doi.org/10.1007/s10072-020-04626-9.

[277] G. Barbagallo, G. Arabia, F. Novellino, R. Nista, M. Salsone, M. Morelli, F. Rocca, A. Quattrone, M. Caracciolo, U. Sabatini, A. Cherubini, A. Quattrone, Increased glutamate plus glutamine levels in the thalamus of patients with essential tremor: A preliminary proton MR spectroscopic study, Parkinsonism Relat. Disord. 47 (2018) 57–63. https://doi.org/10.1016/j.parkreldis.2017.11.345.

[278] I. Cheong, M. Marjanska, D.K. Deelchand, L.E. Eberly, D. Walk, G. Oz, Ultra-High Field Proton MR Spectroscopy in Early-Stage Amyotrophic Lateral Sclerosis, Neurochem. Res. 42 (2017) 1833–1844. https://doi.org/10.1007/s11064-017-2248-2.

[279] A.A. Vijayakumari, R.N. Menon, B. Thomas, T.M. Arun, M. Nandini, C. Kesavadas, Glutamatergic response to a low load working memory paradigm in the left dorsolateral prefrontal cortex in patients with mild cognitive impairment: a functional magnetic resonance spectroscopy study, Brain Imaging Behav. 14 (2020) 451–459. https://doi.org/10.1007/s11682-019-00122-7.

[280] Z. Guo, X. Liu, H. Hou, F. Wei, X. Chen, Y. Shen, W. Chen, 1H-MRS asymmetry changes in the anterior and posterior cingulate gyrus in patients with mild cognitive impairment and mild Alzheimer’s disease, Compr. Psychiatry. 69 (2016) 179–185. https://doi.org/10.1016/j.comppsych.2016.06.001.

[281] O. Voevodskaya, K. Poulakis, P. Sundgren, D. Van Westen, S. Palmqvist, L.O. Wahlund, E. Stomrud, O. Hansson, E. Westman, Brain myoinositol as a potential marker of amyloid-related pathology: A longitudinal study, Neurology. 92 (2019) E395–E405. https://doi.org/10.1212/WNL.0000000000006852.

[282] Y.G. Khomenko, G. V. Kataeva, A.A. Bogdan, E.M. Chernysheva, D.S. Susin, Cerebral Metabolism in Patients with Cognitive Disorders: a Combined Magnetic Resonance Spectroscopy and Positron Emission Tomography Study, Neurosci. Behav. Physiol. 49 (2019) 1199–1207. https://doi.org/10.1007/s11055-019-00858-1.

[283] L. Glodzik, M. Sollberger, A. Gass, A. Gokhale, H. Rusinek, J.S. Babb, J.G. Hirsch, M. Amann, A.U. Monsch, O. Gonen, Global N-acetylaspartate in normal subjects, mild cognitive impairment and Alzheimer’s disease patients, J. Alzheimer’s Dis. 43 (2015) 939–947. https://doi.org/10.3233/JAD-140609.

[284] N. Seraji-Bozorgzad, F. Bao, E. George, S. Krstevska, V. Gorden, J. Chorostecki, C. Santiago, I. Zak, C. Caon, O. Khan, Longitudinal study of the substantia nigra in Parkinson disease: A high-field 1H-MR spectroscopy imaging study, Mov. Disord. 30 (2015) 1400–1404. https://doi.org/10.1002/mds.26323.

[285] E.D. Louis, N. Hernandez, J.P. Dyke, R.E. Ma, U. Dydak, In Vivo Dentate Nucleus Gamma-aminobutyric Acid Concentration in Essential Tremor vs. Controls, Cerebellum. 17 (2018) 165–172. https://doi.org/10.1007/s12311-017-0891-4.

[286] B. Zeydan, D.K. Deelchand, N. Tosakulwong, T.G. Lesnick, O.H. Kantarci, M.M. Machulda, D.S. Knopman, V.J. Lowe, C.R. Jack, R.C. Petersen, G. Oz, K. Kantarci, Decreased Glutamate Levels in Patients with Amnestic Mild Cognitive Impairment: An sLASER Proton MR Spectroscopy and PiB-PET Study, J. NEUROIMAGING. 27 (2017) 630–636. https://doi.org/10.1111/jon.12454.

[287] T. Gong, Y.Y. Xiang, M.G. Saleh, F. Gao, W.B. Chen, R.A.E. Edden, G.B. Wang, Inhibitory Motor Dysfunction in Parkinson’s Disease Subtypes, J. Magn. Reson. IMAGING. 47 (2018) 1610–1615. https://doi.org/10.1002/jmri.25865.

[288] J.J. Gomar, M.L. Gordon, D. Dickinson, P.B. Kingsley, A.M. Ulug, L. Keehlisen, S. Huet, J.J. Buthorn, J. Koppel, E. Christen, C. Conejero-Goldberg, P. Davies, T.E. Goldberg, APOE Genotype Modulates Proton Magnetic Resonance Spectroscopy Metabolites in the Aging Brain, Biol. Psychiatry. 75 (2014) 686–692. https://doi.org/10.1016/j.biopsych.2013.05.022.

[289] M.İ. Atagün, E.M. Şıkoğlu, S.S. Can, G.K. Uğurlu, S.U. Kaymak, A. Çayköylü, O. Algın, M.L. Phillips, C.M. Moore, D. Öngür, Neurochemical differences between bipolar disorder type I and II in superior temporal cortices: A proton magnetic resonance spectroscopy study, J. Affect. Disord. 235 (2018) 15–19. https://doi.org/10.1016/j.jad.2018.04.010.

[290] K.A. Bradley, C.M. Alonso, L.M. Mehra, J.Q. Xu, V. Gabbay, Elevated striatal gamma-aminobutyric acid in youth with major depressive disorder, Prog. Neuropsychopharmacol. Biol. Psychiatry. 86 (2018) 203–210. https://doi.org/10.1016/j.pnpbp.2018.06.004.

[291] C.M. Cirstea, P. Lee, S.C. Craciunas, I.Y. Choi, J.E. Burris, R.J. Nudo, Pre-therapy Neural State of Bilateral Motor and Premotor Cortices Predicts Therapy Gain After Subcortical Stroke: A Pilot Study, Am. J. Phys. Med. Rehabil. 97 (2018) 23–33. https://doi.org/10.1097/PHM.0000000000000791.

[292] J.E. Kim, G.H. Kim, J. Hwang, J.Y. Kim, P.F. Renshaw, D.A. Yurgelun-Todd, B. Kim, I. Kang, S. Jeon, J. Ma, I.K. Lyoo, S. Yoon, Metabolic alterations in the anterior cingulate cortex and related cognitive deficits in late adolescent methamphetamine users, Addict. Biol. 23 (2018) 327–336. https://doi.org/10.1111/adb.12473.

[293] Y. Li, M. Lafontaine, S.S. Chang, S.J. Nelson, Comparison between Short and Long Echo Time Magnetic Resonance Spectroscopic Imaging at 3T and 7T for Evaluating Brain Metabolites in Patients with Glioma, ACS Chem. Neurosci. 9 (2018) 130–137. https://doi.org/10.1021/acschemneuro.7b00286.

[294] C. Sheth, A. Prescot, E. Bueler, J. DiMuzio, M. Legarreta, P.F. Renshaw, D. Yurgelun-Todd, E. McGlade, Alterations in anterior cingulate cortex myoinositol and aggression in veterans with suicidal behavior: A proton magnetic resonance spectroscopy study, Psychiatry Res. - Neuroimaging. 276 (2018) 24–32. https://doi.org/10.1016/j.pscychresns.2018.04.004.

[295] S. Sivaraman, N. V Kraguljac, D.M. White, C.J. Morgan, S.S. Gonzales, A.C. Lahti, Neurometabolic abnormalities in the associative striatum in antipsychotic-naive first episode psychosis patients, PSYCHIATRY Res. 281 (2018) 101–106. https://doi.org/10.1016/j.pscychresns.2018.06.003.

[296] F. Branzoli, C. Pontoizeau, L. Tchara, A.L. Di Stefano, A. Kamoun, D.K. Deelchand, R. Valabrègue, S. Lehéricy, M. Sanson, C. Ottolenghi, M. Marjańska, Cystathionine as a marker for 1p/19q codeleted gliomas by in vivo magnetic resonance spectroscopy, Neuro. Oncol. 21 (2019) 765–774. https://doi.org/10.1093/neuonc/noz031.

[297] N. Fayed, B. Oliván, Y. Lopez del Hoyo, E. Andrés, M.C. Perez-Yus, A. Fayed, L.F. Angel, A. Serrano-Blanco, M. Roca, J. Garcia Campayo, Changes in metabolites in the brain of patients with fibromyalgia after treatment with an NMDA receptor antagonist, Neuroradiol. J. 32 (2019) 408–419. https://doi.org/10.1177/1971400919857544.

[298] R.R. Girgis, S. Baker, X.L. Mao, R. Gil, D.C. Javitt, J.T. Kantrowitz, M. Gu, D.M. Spielman, N. Ojeil, X.Y. Xu, A. Abi-Dargham, D.C. Shungu, L.S. Kegeles, Effects of acute N-acetylcysteine challenge on cortical glutathione and glutamate in schizophrenia: A pilot in vivo proton magnetic resonance spectroscopy study, PSYCHIATRY Res. 275 (2019) 78–85. https://doi.org/10.1016/j.psychres.2019.03.018.

[299] T.M. Hansen, B. Brock, A. Juhl, A.M. Drewes, H. Vorum, C.U. Andersen, P.E. Jakobsen, J. Karmisholt, J.B. Frøkjær, C. Brock, Brain spectroscopy reveals that N-acetylaspartate is associated to peripheral sensorimotor neuropathy in type 1 diabetes, J. Diabetes Complications. 33 (2019) 323–328. https://doi.org/10.1016/j.jdiacomp.2018.12.016.

[300] E.J. Meyer, J.N. Stout, A.W. Chung, P.E. Grant, R. Mannix, B. Gagoski, Longitudinal Changes in Magnetic Resonance Spectroscopy in Pediatric Concussion: A Pilot Study, Front. Neurol. 10 (2019) 1–7. https://doi.org/10.3389/fneur.2019.00556.

[301] D. Didehdar, F. Kamali, A.K. Yoosefinejad, M. Lotfi, The effect of spinal manipulation on brain neurometabolites in chronic nonspecific low back pain patients: a randomized clinical trial, Ir. J. Med. Sci. 189 (2020) 543–550. https://doi.org/10.1007/s11845-019-02140-2.

[302] C.P. Lewis, J.D. Port, C.J. Blacker, A.I. Sonmez, B.J. Seewoo, J.M. Leffler, M.A. Frye, P.E. Croarkin, Altered anterior cingulate glutamatergic metabolism in depressed adolescents with current suicidal ideation, Transl. Psychiatry. 10 (2020). https://doi.org/10.1038/s41398-020-0792-z.

[303] F.P. Macmaster, Q. Mclellan, A.D. Harris, S. Virani, K.M. Barlow, L.M. Langevin, K.O. Yeates, B.L. Brooks, N-Acetyl-Aspartate in the Dorsolateral Prefrontal Cortex Long after Concussion in Youth, J. Head Trauma Rehabil. 35 (2020) E127–E135. https://doi.org/10.1097/HTR.0000000000000535.

[304] N. Mazibuko, R.O.G. Tuura, L. Sztriha, O. O’Daly, G.J. Barker, S.C.R. Williams, M. O’Sullivan, L. Kalra, Subacute changes in N-acetylaspartate (NAA) following Ischemic Stroke: A Serial MR Spectroscopy Pilot Study, Diagnostics. 10 (2020) 1–12. https://doi.org/10.3390/diagnostics10070482.

[305] P. Menshchikov, A. Ivantsova, A. Manzhurtsev, M. Ublinskiy, A. Yakovlev, I. Melnikov, D. Kupriyanov, T. Akhadov, N. Semenova, Separate N-acetyl aspartyl glutamate, N-acetyl aspartate, aspartate, and glutamate quantification after pediatric mild traumatic brain injury in the acute phase, Magn. Reson. Med. 84 (2020) 2918–2931. https://doi.org/10.1002/mrm.28332.

[306] S. Oleson, D. Eagan, S. Kaur, W.J. Hertzing, M. Alkatan, J.N. Davis, H. Tanaka, A.P. Haley, Apolipoprotein E genotype moderates the association between dietary polyunsaturated fat and brain function: an exploration of cerebral glutamate and cognitive performance, Nutr. Neurosci. 23 (2020) 696–705. https://doi.org/10.1080/1028415X.2018.1547857.

[307] S.D. Pizzi, R. Franciotti, A. Ferretti, R.A.E. Edden, H.J. Zollner, R. Esposito, G. Bubbico, C. Aiello, F. Calvanese, S.L. Sensi, A. Tartaro, M. Onofrj, L. Bonanni, High gamma-Aminobutyric Acid Content within the Medial Prefrontal Cortex Is a Functional Signature of Somatic Symptoms Disorder in Patients with Parkinson’s Disease, Mov. Disord. 35 (2020) 2184–2192. https://doi.org/10.1002/mds.28221.

[308] C. Sheth, A.P. Prescot, M. Legarreta, P.F. Renshaw, E. McGlade, D. Yurgelun-Todd, Increased myoinositol in the anterior cingulate cortex of veterans with a history of traumatic brain injury: A proton magnetic resonance spectroscopy study, J. Neurophysiol. 123 (2020) 1619–1629. https://doi.org/10.1152/jn.00765.2019.

[309] C.E. Wiers, S.I. Cunningham, D.G. Tomasi, T. Ernst, L. Chang, E. Shokri-Kojori, G.J. Wang, N.D. Volkow, Elevated thalamic glutamate levels and reduced water diffusivity in alcohol use disorder: Association with impulsivity, PSYCHIATRY Res. 305 (2020). https://doi.org/10.1016/j.pscychresns.2020.111185.

[310] D. Wong, S. Atiya, J. Fogarty, M. Montero-Odasso, S.H. Pasternak, C. Brymer, M.J. Borrie, R. Bartha, Reduced Hippocampal Glutamate and Posterior Cingulate N-Acetyl Aspartate in Mild Cognitive Impairment and Alzheimer’s Disease Is Associated with Episodic Memory Performance and White Matter Integrity in the Cingulum: A Pilot Study, J. Alzheimer’s Dis. 73 (2020) 1385–1405. https://doi.org/10.3233/JAD-190773.

[311] H. Zheng, W. Yang, B. Zhang, G. Hua, S. Wang, F. Jia, G. Guo, W. Wang, D. Quan, Reduced anterior cingulate glutamate of comorbid skin-picking disorder in adults with obsessive-compulsive disorder, J. Affect. Disord. 265 (2020) 193–199. https://doi.org/10.1016/j.jad.2020.01.059.

[312] I. Mawla, E. Ichesco, H.J. Zöllner, R.A.E. Edden, T. Chenevert, H. Buchtel, M.D. Bretz, H. Sloan, C.M. Kaplan, S.E. Harte, G.A. Mashour, D.J. Clauw, V. Napadow, R.E. Harris, Greater Somatosensory Afference With Acupuncture Increases Primary Somatosensory Connectivity and Alleviates Fibromyalgia Pain via Insular γ-Aminobutyric Acid: A Randomized Neuroimaging Trial, Arthritis Rheumatol. 73 (2021) 1318–1328. https://doi.org/10.1002/art.41620.

[313] A. O’Neill, L. Annibale, G. Blest-Hopley, R. Wilson, V. Giampietro, S. Bhattacharyya, Cannabidiol modulation of hippocampal glutamate in early psychosis, J. Psychopharmacol. 35 (2021) 814–822. https://doi.org/10.1177/02698811211001107.

[314] J.O. Friedrich, N.K.J. Adhikari, J. Beyene, Ratio of means for analyzing continuous outcomes in meta-analysis performed as well as mean difference methods, J. Clin. Epidemiol. 64 (2011) 556–564. https://doi.org/10.1016/j.jclinepi.2010.09.016.

[315] J.O. Friedrich, N.K.J. Adhikari, J. Beyene, The ratio of means method as an alternative to mean differences for analyzing continuous outcome variables in meta-analysis: A simulation study, BMC Med. Res. Methodol. 8 (2008). https://doi.org/10.1186/1471-2288-8-32.

[316] R. Longo, A. Bampo, R. Vidimari, S. Magnaldi, A. Giorgini, Absolute Quantitation of Brain 1H Nuclear Magnetic Resonance Spectra, Invest. Radiol. 30 (1995) 199–203. https://doi.org/10.1097/00004424-199504000-00001.

[317] S. Posse, C.A. Cuenod, R. Risinger, D. Le Bihan, R.S. Balaban, Anomalous Transverse Relaxation in 1H Spectroscopy in Human Brain at 4 Tesla, Magn. Reson. Med. 33 (1995) 246–252. https://doi.org/10.1002/mrm.1910330215.

[318] A. Van Der Toorn, R.M. Dijkhuizen, C.A.F. Tulleken, K. Nicolay, T1 and T2 relaxation times of the major 1H-containing metabolites in rat brain after focal ischemia, NMR Biomed. 8 (1995) 245–252. https://doi.org/10.1002/nbm.1940080604.

[319] E.B. Cady, Metabolite concentrations and relaxation in perinatal cerebral hypoxic-ischemic injury, Neurochem. Res. 21 (1996) 1043–1052. https://doi.org/10.1007/BF02532414.

[320] E.B. Cady, J. Penrice, P.N. Amess, A. Lorek, M. Wylezinska, R.F. Aldridge, F. Franconi, J.S. Wyatt, E.O.R. Reynolds, Lactate,N-acetylaspartate, choline and creatine concentrations, and spin-spin relaxation in thalamic and occipito-parietal regions of developing human brain, Magn. Reson. Med. 36 (1996) 878–886. https://doi.org/10.1002/mrm.1910360610.

[321] W. Block, J. Karitzky, F. Träber, C. Pohl, E. Keller, R.R. Mundegar, R. Lamerichs, H. Rink, F. Ries, H.H. Schild, F. Jerusalem, Proton magnetic resonance spectroscopy of the primary motor cortex in patients with motor neuron disease: Subgroup analysis and follow-up measurements, Arch. Neurol. 55 (1998) 931–936. https://doi.org/10.1001/archneur.55.7.931.

[322] H. Fujimori, T. Michaelis, M. Wick, J. Frahm, Proton T2 relaxation of cerebral metabolites during transient global ischemia in rat brain, Magn. Reson. Med. 39 (1998) 647–650. https://doi.org/10.1002/mrm.1910390419.

[323] C.G. Choi, J. Frahm, Localized proton MRS of the human hippocampus: Metabolite concentrations and relaxation times, Magn. Reson. Med. 41 (1999) 204–207. https://doi.org/10.1002/(SICI)1522-2594(199901)41:1<204::AID-MRM29>3.0.CO;2-7.

[324] P.B. Barker, D.O. Hearshen, M.D. Boska, Single-voxel proton MRS of the human brain at 1.5T and 3.0T, Magn. Reson. Med. 45 (2001) 765–769. https://doi.org/10.1002/mrm.1104.

[325] J.C.W. Brooks, N. Roberts, G.J. Kemp, M.A. Gosney, M. Lye, G.H. Whitehouse, A Proton Magnetic Resonance Spectroscopy Study of Age-related Changes in Frontal Lobe Metabolite Concentrations, Cereb. Cortex. 11 (2001) 598–605. https://doi.org/10.1093/cercor/11.7.598.

[326] P.A. Narayana, D. Johnston, D.P. Flamig, In vivo proton magnetic resonance spectroscopy studies of human brain, Magn. Reson. Imaging. 9 (1991) 303–308. https://doi.org/10.1016/0730-725X(91)90415-I.

[327] I. Tkáč, P. Andersen, G. Adriany, H. Merkle, K. Uǧurbil, R. Gruetter, In vivo 1 H NMR spectroscopy of the human brain at 7 T, Magn. Reson. Med. 46 (2001) 451–456. https://doi.org/10.1002/mrm.1213.

[328] C.C. Hanstock, V.A. Cwik, W.R.W. Martin, Reduction in metabolite transverse relaxation times in amyotrophic lateral sclerosis, J. Neurol. Sci. 198 (2002) 37–41. https://doi.org/10.1016/S0022-510X(02)00074-6.

[329] M. Mascalchi, R. Brugnoli, L. Guerrini, G. Belli, M. Nistri, L.S. Politi, C. Gavazzi, F. Lolli, G. Argenti, N. Villari, Single-voxel long TE 1H-MR spectroscopy of the normal brainstem and cerebellum, J. Magn. Reson. Imaging. 16 (2002) 532–537. https://doi.org/10.1002/jmri.10189.

[330] S. Michaeli, M. Garwood, X.-H. Zhu, L. DelaBarre, P. Andersen, G. Adriany, H. Merkle, K. Ugurbil, W. Chen, ProtonT2 relaxation study of water, N-acetylaspartate, and creatine in human brain using Hahn and Carr-Purcell spin echoes at 4T and 7T, Magn. Reson. Med. 47 (2002) 629–633. https://doi.org/10.1002/mrm.10135.

[331] D.R. Rutgers, J. van der Grond, Relaxation times of choline, creatine andN-acetyl aspartate in human cerebral white matter at 1.5 T, NMR Biomed. 15 (2002) 215–221. https://doi.org/10.1002/nbm.762.

[332] P. Sarchielli, O. Presciutti, R. Tarducci, G. Gobbi, A. Alberti, G.P. Pelliccioli, P. Chiarini, V. Gallai, Localized 1H magnetic resonance spectroscopy in mainly cortical gray matter of patients with multiple sclerosis, J. Neurol. 249 (2002) 902–910. https://doi.org/10.1007/s00415-002-0758-5.

[333] P.E. Sijens, M. Oudkerk, 1H chemical shift imaging characterization of human brain tumor and edema, Eur. Radiol. 12 (2002) 2056–2061. https://doi.org/10.1007/s00330-001-1300-3.

[334] H. Lei, Y. Zhang, X.H. Zhu, W. Chen, Changes in the proton T2 relaxation times of cerebral water and metabolites during forebrain ischemia in rat at 9.4 T, Magn. Reson. Med. 49 (2003) 979–984. https://doi.org/10.1002/mrm.10490.

[335] R. Hurd, N. Sailasuta, R. Srinivasan, D.B. Vigneron, D. Pelletier, S.J. Nelson, Measurement of brain glutamate using TE-averaged PRESS at 3T, Magn. Reson. Med. 51 (2004) 435–440. https://doi.org/10.1002/mrm.20007.

[336] F. Träber, W. Block, R. Lamerichs, J. Gieseke, H.H. Schild, 1H Metabolite Relaxation Times at 3.0 Tesla: Measurements of T1 and T2 Values in Normal Brain and Determination of Regional Differences in Transverse Relaxation, J. Magn. Reson. Imaging. 19 (2004) 537–545. https://doi.org/10.1002/jmri.20053.

[337] P. Gideon, O. Henriksen, In vivo relaxation of N-acetyl-aspartate, creatine plus phosphocreatine, and choline containing compounds during the course of brain infarction: A proton MRS study, Magn. Reson. Imaging. 10 (1992) 983–988. https://doi.org/10.1016/0730-725X(92)90453-7.

[338] P.M. Walker, D. Ben Salem, A. Lalande, M. Giroud, F. Brunotte, Time course of NAA T2 and ADCw in ischaemic stroke patients: 1H MRS imaging and diffusion-weighted MRI, J. Neurol. Sci. 220 (2004) 23–28. https://doi.org/10.1016/j.jns.2004.01.012.

[339] E.E. Brief, K.P. Whittall, D.K.B. Li, A.L. MacKay, Proton T2 relaxation of cerebral metabolites of normal human brain over large TE range, NMR Biomed. 18 (2005) 14–18. https://doi.org/10.1002/nbm.916.

[340] M. Dumoulin, E. Zimmerman, R. Hurd, I. Hancu, Increased brain metabolite T 2 relaxation times in patients with Alzheimer’s disease, Proc. 13-Th Meet. Int. Soc. Magn. Reson. Med. (2005) 1179.

[341] R. Kreis, J. Slotboom, L. Hofmann, C. Boesch, Integrated data acquisition and processing to determine metabolite contents, relaxation times, and macromolecule baseline in single examinations of individual subjects, Magn. Reson. Med. 54 (2005) 761–768. https://doi.org/10.1002/mrm.20673.

[342] B.J. Soher, P.M. Pattany, G.B. Matson, A.A. Maudsley, Observation of coupled 1H metabolite resonances at long TE, Magn. Reson. Med. 53 (2005) 1283–1287. https://doi.org/10.1002/mrm.20491.

[343] C. Choi, N.J. Coupland, P.P. Bhardwaj, S. Kalra, C.A. Casault, K. Reid, P.S. Allen, T2 measurement and quantification of glutamate in human brain in vivo, Magn. Reson. Med. 56 (2006) 971–977. https://doi.org/10.1002/mrm.21055.

[344] W. Zaaraoui, L. Fleysher, R. Fleysher, S. Liu, B.J. Soher, O. Gonen, Human brain-structure resolved T2 relaxation times of proton metabolites at 3 Tesla, Magn. Reson. Med. 57 (2007) 983–989. https://doi.org/10.1002/mrm.21250.

[345] C. Chassain, G. Bielicki, E. Durand, S. Lolignier, F. Essafi, A. Traoré, F. Durif, Metabolic changes detected by proton magnetic resonance spectroscopy in vivo and in vitro in a murin model of Parkinson’s disease, the MPTP-intoxicated mouse, J. Neurochem. 105 (2008) 874– 882. https://doi.org/10.1111/j.1471-4159.2007.05185.x.

[346] C. Cudalbu, A. Rengle, O. Beuf, S. Cavassila, Rat brain metabolite relaxation time estimates using magnetic resonance spectroscopy at two different field strengths, Comptes Rendus Chim. 11 (2008) 442–447. https://doi.org/10.1016/j.crci.2007.08.013.

[347] I.I. Kirov, L. Fleysher, R. Fleysher, V. Patil, S. Liu, O. Gonen, Age dependence of regional proton metabolites T2 relaxation times in the human brain at 3 T, Magn. Reson. Med. 60 (2008) 790–795. https://doi.org/10.1002/mrm.21715.

[348] J. Hennig, H. Pfister, T. Ernst, D. Ott, Direct absolute quantification of metabolites in the human brain with in vivo localized proton spectroscopy, NMR Biomed. 5 (1992) 193–199. https://doi.org/10.1002/nbm.1940050406.

[349] Y. Li, R. Srinivasan, H. Ratiney, Y. Lu, S.M. Chang, S.J. Nelson, Comparison of T1 and T2 metabolite relaxation times in glioma and normal brain at 3T, J. Magn. Reson. Imaging. 28 (2008) 342–350. https://doi.org/10.1002/jmri.21453.

[350] N. Tunc-Skarka, W. Weber-Fahr, M. Hoerst, A. Meyer-Lindenberg, M. Zink, G. Ende, MR spectroscopic evaluation of N-acetylaspartate’s T2 relaxation time and concentration corroborates white matter abnormalities in schizophrenia, Neuroimage. 48 (2009) 525–531. https://doi.org/10.1016/j.neuroimage.2009.06.061.

[351] D.K. Deelchand, P.-F. Van de Moortele, G. Adriany, I. Iltis, P. Andersen, J.P. Strupp, J. Thomas Vaughan, K. Uğurbil, P.-G. Henry, In vivo 1H NMR spectroscopy of the human brain at 9.4T: Initial results, J. Magn. Reson. 206 (2010) 74–80. https://doi.org/10.1016/j.jmr.2010.06.006.

[352] U.E. Emir, D. Deelchand, P.G. Henry, M. Terpstra, Noninvasive quantification of T2 and concentrations of ascorbate and glutathione in the human brain from the same double-edited spectra, NMR Biomed. 24 (2011) 263–269. https://doi.org/10.1002/nbm.1583.

[353] M. Marjańska, E.J. Auerbach, R. Valabrègue, P.F. Van de Moortele, G. Adriany, M. Garwood, Localized 1H NMR spectroscopy in different regions of human brain in vivo at 7T: T 2 relaxation times and concentrations of cerebral metabolites, NMR Biomed. 25 (2012) 332–339. https://doi.org/10.1002/nbm.1754.

[354] D.K. Deelchand, P.G. Henry, K. Uǧurbil, M. Marjańska, Measurement of transverse relaxation times of J-coupled metabolites in the human visual cortex at 4 T, Magn. Reson. Med. 67 (2012) 891–897. https://doi.org/10.1002/mrm.23080.

[355] F. Du, A. Cooper, B.M. Cohen, P.F. Renshaw, D. Öngür, Water and metabolite transverse T2 relaxation time abnormalities in the white matter in schizophrenia, Schizophr. Res. 137 (2012) 241–245. https://doi.org/10.1016/j.schres.2012.01.026.

[356] R.A.E. Edden, J. Intrapiromkul, H. Zhu, Y. Cheng, P.B. Barker, Measuring T 2 in vivo with J-difference editing: Application to GABA at 3 tesla, J. Magn. Reson. Imaging. 35 (2012) 229–234. https://doi.org/10.1002/jmri.22865.

[357] S.K. Ganji, A. Banerjee, A.M. Patel, Y.D. Zhao, I.E. Dimitrov, J.D. Browning, E. Sherwood Brown, E.A. Maher, C. Choi, T 2 measurement of J-coupled metabolites in the human brain at 3T, NMR Biomed. 25 (2012) 523–529. https://doi.org/10.1002/nbm.1767.

[358] Y. Li, T1 and T2 Metabolite Relaxation Times in Normal Brain at 3T and 7T, J. Mol. Imaging Dyn. 02 (2013) 1–5. https://doi.org/10.4172/2155-9937.s1-002.

[359] P. Christiansen, P. Toft, H.B.W. Larsson, M. Stubgaard, O. Henriksen, The concentration of N-acetyl aspartate, creatine + phosphocreatine, and choline in different parts of the brain in adulthood and senium, Magn. Reson. Imaging. 11 (1993) 799–806. https://doi.org/10.1016/0730-725X(93)90197-L.

[360] A. Andreychenko, D.W.J. Klomp, R.A. De Graaf, P.R. Luijten, V.O. Boer, In vivo GABA T2 determination with J-refocused echo time extension at 7 T, NMR Biomed. 26 (2013) 1596– 1601. https://doi.org/10.1002/nbm.2997.

[361] M. Marjańska, U.E. Emir, D.K. Deelchand, M. Terpstra, Faster Metabolite 1H Transverse Relaxation in the Elder Human Brain, PLoS One. 8 (2013) 1–7. https://doi.org/10.1371/journal.pone.0077572.

[362] I. Ronen, E. Ercan, A. Webb, Rapid multi-echo measurement of brain metabolite T2 values at 7T using a single-shot spectroscopic Carr-Purcell-Meiboom-Gill sequence and prior information, NMR Biomed. 26 (2013) 1291–1298. https://doi.org/10.1002/nbm.2951.

[363] F. Branzoli, E. Ercan, A. Webb, I. Ronen, The interaction between apparent diffusion coefficients and transverse relaxation rates of human brain metabolites and water studied by diffusion-weighted spectroscopy at 7 T, NMR Biomed. 27 (2014) 495–506. https://doi.org/10.1002/nbm.3085.

[364] A.P. Prescot, X. Shi, C. Choi, P.F. Renshaw, In vivo T 2 relaxation time measurement with echo-time averaging, NMR Biomed. 27 (2014) 863–869. https://doi.org/10.1002/nbm.3115.

[365] D.K. Deelchand, P.G. Henry, M. Marjańska, Effect of Carr-Purcell refocusing pulse trains on transverse relaxation times of metabolites in rat brain at 9.4 Tesla, Magn. Reson. Med. 73 (2015) 13–20. https://doi.org/10.1002/mrm.25088.

[366] A. Madan, S.K. Ganji, Z. An, K.S. Choe, M.C. Pinho, R.M. Bachoo, E.M. Maher, C. Choi, Proton T2 measurement and quantification of lactate in brain tumors by MRS at 3 Tesla in vivo, Magn. Reson. Med. 73 (2015) 2094–2099. https://doi.org/10.1002/mrm.25352.

[367] Y. Zhang, J. Shen, Simultaneous quantification of glutamate and glutamine by J-modulated spectroscopy at 3 Tesla, Magn. Reson. Med. 76 (2016) 725–732. https://doi.org/10.1002/mrm.25922.

[368] M.E. Fisher, B.J. Dobberthien, A.G. Tessier, A. Yahya, Characterization of the response of taurine protons to PRESS at 9.4 T for Resolving choline and Determining taurine T2, NMR Biomed. 29 (2016) 1427–1435. https://doi.org/10.1002/nbm.3588.

[369] F. Jiru, A. Skoch, D. Wagnerova, M. Dezortova, J. Viskova, O. Profant, J. Syka, M. Hajek, The age dependence of T2 relaxation times of N-acetyl aspartate, creatine and choline in the human brain at 3 and 4T, NMR Biomed. 29 (2016) 284–292. https://doi.org/10.1002/nbm.3456.

[370] R. Kreis, T. Ernst, B.D. Ross, Development of the human brain: In vivo quantification of metabolite and water content with proton magnetic resonance spectroscopy, Magn. Reson. Med. 30 (1993) 424–437. https://doi.org/10.1002/mrm.1910300405.

[371] L. An, S. Li, J. Shen, Simultaneous determination of metabolite concentrations, T1 and T2 relaxation times, Magn. Reson. Med. 78 (2017) 2072–2081. https://doi.org/10.1002/mrm.26612.

[372] C. Chen, B. Lanz, C. Fernandes, S. Francis, P. Gowland, P. Morris, The use of MEGA-sLASER with J-refocusing echo time extension to measure the proton T2 of lactate in healthy human brain at 7 T, Proceeding Int. Soc. Magn. Reson. Med. (2017) 5494.

[373] D.K. Deelchand, E.J. Auerbach, N. Kobayashi, M. Marjańska, Transverse relaxation time constants of the five major metabolites in human brain measured in vivo using LASER and PRESS at 3 T, Magn. Reson. Med. 79 (2018) 1260–1265. https://doi.org/10.1002/mrm.26826.

[374] L. Li, N. Li, L. An, J. Shen, A novel approach to probing in vivo metabolite relaxation: Linear quantification of spatially modulated magnetization, Magn. Reson. Med. 79 (2018) 2491–2499. https://doi.org/10.1002/mrm.26941.

[375] K.M. Swanberg, H. Prinsen, D. Coman, R.A. de Graaf, C. Juchem, Quantification of glutathione transverse relaxation time T2 using echo time extension with variable refocusing selectivity and symmetry in the human brain at 7 Tesla, J. Magn. Reson. 290 (2018) 1–11. https://doi.org/10.1016/j.jmr.2018.02.017.

[376] D. Wong, A.L. Schranz, R. Bartha, Optimized in vivo brain glutamate measurement using long-echo-time semi-LASER at 7 T, NMR Biomed. 31 (2018) 1–13. https://doi.org/10.1002/nbm.4002.

[377] P.O. Wyss, C. Bianchini, M. Scheidegger, I.A. Giapitzakis, A. Hock, A. Fuchs, A. Henning, In vivo estimation of transverse relaxation time constant (T2) of 17 human brain metabolites at 3T, Magn. Reson. Med. 80 (2018) 452–461. https://doi.org/10.1002/mrm.27067.

[378] N. Li, L. Li, Y. Zhang, M.F. Araneta, C. Johnson, J. Shen, Quantification of in vivo transverse relaxation of glutamate in the frontal cortex of human brain by radio frequency pulse-driven longitudinal steady state, PLoS One. 14 (2019) 1–13. https://doi.org/10.1371/journal.pone.0215210.

[379] P. Menshchikov, A. Manzhurtsev, M. Ublinskiy, T. Akhadov, N. Semenova, T2measurement and quantification of cerebral white and gray matter aspartate concentrations in vivo at 3T: a MEGA-PRESS study, Magn. Reson. Med. 82 (2019) 11–20. https://doi.org/10.1002/mrm.27700.

[380] D.K. Deelchand, J.R. McCarten, L.S. Hemmy, E.J. Auerbach, L.E. Eberly, M. Marjańska, Changes in the intracellular microenvironment in the aging human brain, Neurobiol. Aging. 95 (2020) 168–175. https://doi.org/10.1016/j.neurobiolaging.2020.07.017.

[381] A.M. Blamire, G.D. Graham, D.L. Rothman, J.W. Prichard, Proton spectroscopy of human stroke: Assessment of transverse relaxation times and partial volume effects in single volume STEAM MRS, Magn. Reson. Imaging. 12 (1994) 1227–1235. https://doi.org/10.1016/0730-725X(94)90087-8.

[382] S. Murali-Manohar, T. Borbath, A.M. Wright, B. Soher, R. Mekle, A. Henning, T2 relaxation times of macromolecules and metabolites in the human brain at 9.4 T, Magn. Reson. Med. 84 (2020) 542–558. https://doi.org/10.1002/mrm.28174.

[383] C.-H. Yoo, H.-M. Baek, K.-H. Song, D.-C. Woo, B.-Y. Choe, An in vivo proton magnetic resonance spectroscopy study with optimized echo-time technique for concurrent quantification and T2 measurement targeting glutamate in the rat brain, Magn. Reson. Mater. Physics, Biol. Med. 33 (2020) 735–746. https://doi.org/10.1007/s10334-020-00840-w.

[384] M. Dacko, T. Lange, Flexible MEGA editing scheme with asymmetric adiabatic pulses applied for measurement of lactate in human brain, Magn. Reson. Med. 85 (2021) 1160–1174. https://doi.org/10.1002/mrm.28500.

[385] R. Nosrati, M. Balasubramanian, R. Mulkern, Measuring transverse relaxation rates of the major brain metabolites from single-voxel PRESS acquisitions at a single TE, Magn. Reson. Med. 85 (2021) 2965–2977. https://doi.org/10.1002/mrm.28644.

[386] X. Wang, R. Li, R. He, F. Fang, Effects of repeated manganese treatment on proton magnetic resonance spectra of the globus pallidus in rat brain, NMR Biomed. 35 (2022) 1–10. https://doi.org/10.1002/nbm.4617.

[387] X. Chen, X. Fan, X. Song, M. Gardner, F. Du, D. Öngür, White Matter Metabolite Relaxation and Diffusion Abnormalities in First-Episode Psychosis: A Longitudinal Study, Schizophr. Bull. (2022) 1–9. https://doi.org/10.1093/schbul/sbab149.

[388] K. Kamada, K. Houkin, K. Hida, H. Matsuzawa, Y. Iwasaki, H. Abe, T. Nakada, Localized proton spectroscopy of focal brain pathology in humans: Significant effects of edema on spin– spin relaxation time, Magn. Reson. Med. 31 (1994) 537–540. https://doi.org/10.1002/mrm.1910310510.

[389] P. Christiansen, A. Schlosser, O. Henriksen, Reduced N-acetylaspartate content in the frontal part of the brain in patients with probable Alzheimer’s disease, Magn. Reson. Imaging. 13 (1995) 457–462. https://doi.org/10.1016/0730-725X(94)00113-H.

[390] V. Govindaraju, K. Young, A.A. Maudsley, Proton NMR chemical shifts and coupling constants for brain metabolites, NMR Biomed. 13 (2000) 129–153. https://doi.org/10.1002/1099-1492(200005)13:3<129::AID-NBM619>3.0.CO;2-V.

[391] R.A. de Graaf, In Vivo NMR Spectroscopy, John Wiley & Sons, Ltd, Chichester, UK, 2019. https://doi.org/10.1002/9781119382461.

[392] C. Cleeland, A. Pipingas, A. Scholey, D. White, Neurochemical changes in the aging brain: A systematic review, Neurosci. Biobehav. Rev. 98 (2019) 306–319. https://doi.org/10.1016/j.neubiorev.2019.01.003.

[393] K. Kantarci, C.R. Jack, Y.C. Xu, N.G. Campeau, P.C. O’Brien, G.E. Smith, R.J. Ivnik, B.F. Boeve, E. Kokmen, E.G. Tangalos, R.C. Petersen, Regional metabolic patterns in mild cognitive impairment and Alzheimer’s disease - A H-1 MRS study, Neurology. 55 (2000) 210–217. https://doi.org/10.1212/WNL.55.2.210.

[394] N. Zhang, X. Song, R. Bartha, S. Beyea, R. D’Arcy, Y. Zhang, K. Rockwood, Advances in High-Field Magnetic Resonance Spectroscopy in Alzheimer’s Disease, Curr. Alzheimer Res. 11 (2014) 367–388. https://doi.org/10.2174/1567205011666140302200312.

[395] I. Buard, N. Lopez-Esquibel, F.J. Carey, M.S. Brown, L.D. Medina, E. Kronberg, C.S. Martin, S. Rogers, S.K. Holden, M.R. Greher, B.M. Kluger, Does Prefrontal Glutamate Index Cognitive Changes in Parkinson’s Disease?, Front. Hum. Neurosci. 16 (2022) 1–9. https://doi.org/10.3389/fnhum.2022.809905.

[396] A. Handforth, E.J. Lang, Increased Purkinje Cell Complex Spike and Deep Cerebellar Nucleus Synchrony as a Potential Basis for Syndromic Essential Tremor. A Review and Synthesis of the Literature, Cerebellum. 20 (2021) 266–281. https://doi.org/10.1007/s12311-020-01197-5.

[397] A.T.K. Kendi, F.U. Tan, M. Kendi, H.H. Erdal, S. Tellioǧlu, Magnetic resonance spectroscopy of the thalamus in essential tremor patients, J. Neuroimaging. 15 (2005) 362–366. https://doi.org/10.1177/1051228405279039.

[398] A. Thomson, D. Pasanta, H. Hwa, T. Arichi, R. Edden, X. Chai, N. Puts, No Characterising the lifespan trajectory of six essential neurometabolites in a cohort of 100 participants, in: P. Jezzard (Ed.), Proceeding Int. Soc. Magn. Reson. Med., Toronto, 2023.

[399] T. Gong, S.C.N. Hui, H.J. Zöllner, M. Britton, Y. Song, Y. Chen, A.T. Gudmundson, K.E. Hupfeld, C.W. Davies-Jenkins, S. Murali-Manohar, E.C. Porges, G. Oeltzschner, W. Chen, G. Wang, R.A.E. Edden, Neurometabolic timecourse of healthy aging, Neuroimage. 264 (2022) 119740. https://doi.org/10.1016/j.neuroimage.2022.119740.

[400] P.A. Hardy, D. Gash, R. Yokel, A. Andersen, Y. Ai, Z. Zhang, Correlation of R2 with total iron concentration in the brains of rhesus monkeys, J. Magn. Reson. Imaging. 21 (2005) 118–127. https://doi.org/10.1002/jmri.20244.

[401] C. Langkammer, N. Krebs, W. Goessler, E. Scheurer, F. Ebner, K. Yen, F. Fazekas, S. Ropele, Quantitative MR imaging of brain iron: A postmortem validation study, Radiology. 257 (2010) 455–462. https://doi.org/10.1148/radiol.10100495.

[402] R.J. Ordidge, J.M. Gorell, J.C. Deniau, R.A. Knight, J.A. Helpern, Assessment of relative brain iron concentrations usingT2-weighted andT2*-weighted MRI at 3 Tesla, Magn. Reson. Med. 32 (1994) 335–341. https://doi.org/10.1002/mrm.1910320309.

