## Supplementary Table 1 for "Meta-analysis and Open-source Database for In Vivo Brain Magnetic Resonance Spectroscopy in Health and Disease"

### Supplementary Information:

| **Study** | **Database** | **Search Query** |
| --- | --- | --- |
| Concentration | PubMed | ((Magnetic Resonance Spectroscopy [mh] OR “Magnetic Resonance Spectroscopy” OR MRS OR MRSI OR “^1^H-MRS” OR “In Vivo NMR” OR “In-Vivo NMR” OR “MR Spectroscopy” OR “MRSpectroscopy”) AND (Brain [mh] OR Neurosciences [mh] OR Neurology [mh] OR Neuroscience OR Neurosciences OR Brain OR Midbrain OR Neurology OR Cognition OR Cortical OR Hippocampus OR “Frontal Lobe” OR “Parietal Lobe” OR “Occipital Lobe” OR “Temporal Lobe” OR Amygdala OR Cortex OR Cerebellum OR Cerebrum OR “Brain Stem”) AND (Humans [mh] OR Human OR Humans) AND (Proton OR Protons OR ^1^H OR “Hydrogen Nucleus” OR “^1^H-MRS” OR “^1^H MRS”) NOT (Review)) |
| Concentration | Web of Science | ALL=((“Magnetic Resonance Spectroscopy” OR MRS OR MRSI OR “^1^H-MRS” OR “In Vivo NMR” OR “In-Vivo NMR” OR “MR Spectroscopy” OR “mrspectroscopy”) AND (Neuroscience OR Neurosciences OR Brain OR Midbrain OR Neurology OR Neuroradiology OR Cognition OR Cortical OR Hippocampus OR “Frontal Lobe” OR “Parietal Lobe” OR “Occipital Lobe” OR “Temporal Lobe” OR Amygdala OR Cortex OR Cerebellum OR Cerebrum OR “Brain Stem”) AND (Human OR Humans) AND (Proton OR Protons OR ^1^H OR “Hydrogen Nucleus” OR “^1^H-MRS” OR “^1^H MRS”) NOT (Review)) |
| Concentration | Scopus | (((“magnetic resonance spectroscopy” OR “mrs” OR “^1^H-mrs” OR “mrsi” OR “NMR Spectroscopy”) AND (“human”) AND (“^1^H” OR “proton” OR “^1^H-mrs” OR “hydrogen nucleus”) AND (“neuroscience” OR “brain”) AND (“concentration”) AND NOT (“rat” OR “mouse” OR “mice” OR “rodent” OR “Animal” OR “Dog” OR “ex vivo” OR “ex-vivo” OR “vitro” OR “13C” OR “31P” OR “23Na” OR “17O” OR “Systematic Review”))) |
| Relaxation | PubMed | (“T_2_ Relaxation” OR “Transverse Relaxation” OR “Spin-spin Relaxation” OR “Carr-Purcell Meiboom-Gill”) AND (“Magnetic Resonance Spectroscopy” [mh] OR “NMR Spectroscopy” OR “Magnetic Resonance Spectroscopy” OR “In-Vivo Spectroscopy” OR “Ex-Vivo Spectroscopy”) AND (Brain [mh] OR Neurosciences [mh] OR Neurology [mh] OR “Brain” OR “Neuroscience” OR “Neurology” OR “Phantom”) NOT (“Fingerprinting” OR “CEST” OR “Chemical Exchange Saturation Transfer” OR “31P” OR “(31)P” OR “13C” OR “(13)C” OR “23Na” OR “(23)Na” OR “17O” OR “15N” OR “14N” OR “19F” OR “(19)F” OR “Systematic Review” OR “Food Storage” [mh]) |
| Relaxation | Web of Science | ALL=((“NMR Spectroscopy” OR “Magnetic Resonance Spectroscopy” OR “In-Vivo Spectroscopy” OR “Ex-Vivo Spectroscopy”) AND (Neuroscience OR Neurosciences OR Brain OR Neurology OR Phantom) AND (“T_2_ Relaxation” OR “Transverse Relaxation” OR “Spin-spin Relaxation” OR “Carr-Purcell Meiboom-Gill”) NOT (“Fingerprinting” OR “CEST” OR “Chemical Exchange Saturation Transfer” OR “31P” OR “(31)P” OR “13C” OR “(13)C” OR “23Na” OR “(23)Na” OR “17O” OR “15N” OR “14N” OR “19F” OR “(19)F” OR “Systematic Review” OR “Food Storage”)) |
| Relaxation | Scopus | (((“magnetic resonance spectroscopy” OR “mrs” OR “^1^H-mrs” OR “mrsi” OR “NMR Spectroscopy”) AND (“human”) AND (“^1^H” OR “proton” OR “^1^H-mrs” OR “hydrogen nucleus”) AND (“neuroscience” OR “brain”) AND (“T_2_ Relaxation” OR “Transverse Relaxation” OR “Spin-spin Relaxation” OR “Carr-Purcell Meiboom-Gill”) AND NOT (“13C” OR “31P” OR “23Na” OR “17O” OR “Systematic Review”))) |

**Supplementary Table 1.** Search queries used for each of the databases (PubMed, Web of Science, Scopus) for the Concentration and Relaxation studies.
